# Human-specific tandem repeat in *CACNA1C* modulates responses to neuronal stimulation

**DOI:** 10.1101/2025.09.15.676436

**Authors:** Janet H.T. Song, Fikri Birey, Tzu-Chiao Hung, Nicola A.L. Hall, Catherine A. Guenther, Xiaoyu Chen, Ibrahim F. Alkuraya, Elizabeth M. Tunbridge, Wilfried Haerty, Sergiu P. Pasca, David M. Kingsley

**Affiliations:** Department of Developmental Biology, Stanford University School of Medicine, Stanford, CA, USA; Department of Genetics, Stanford University School of Medicine, Stanford, CA, USA; Department of Psychiatry and Behavioral Sciences, Stanford University, Stanford, CA, USA; Stanford Brain Organogenesis Program, Wu Tsai Neurosciences Institute & Bio-X, Stanford University, Stanford, CA, USA; Department of Psychiatry, University of Oxford, Oxford, UK; Howard Hughes Medical Institute, Stanford University School of Medicine, Stanford, CA, USA; Harvard College, Cambridge, MA, USA; Earlham Institute, Norwich, UK; School of Biology, University of East Anglia, Norwich, UK

## Abstract

The recent development of long-read sequencing has made it possible to catalog variable number tandem repeats (VNTRs) in the human genome. However, little is known about their functional consequences. Here, we characterized the effect of TRACT, a VNTR that is unique to humans and that has sequence variants linked to risk for bipolar disorder and schizophrenia. By adding or removing this VNTR in both mouse models and human neural organoids, we find that TRACT, which is intronic to the L-type voltage-gated calcium channel gene *CACNA1C*, increases intracellular calcium after neuronal stimulation and leads to widespread changes in activity-dependent transcription programs in neurons. TRACT-dependent changes are enriched for genes associated with synapse formation and plasticity, and partially recapitulate evolutionary changes in activity-dependent transcription between species. These findings demonstrate that a single, human-specific, non-coding element can strongly affect the neuronal response to stimulation, and motivate the study of VNTRs as a genetic source of phenotypic variation in both evolution and disease.

## Introduction

Identifying the genetic variants that underlie human evolution and disease risk has proved tremendously challenging. Prior research has used comparative genomics to identify both protein-coding and non-coding sequences that differ between humans and other mammals, display signatures of selection, co-vary with organismal phenotypes, and may contribute to human-specific traits (Pollard et al., 2006; Prabhakar et al., 2008; McLean et al., 2011; Keough et al., 2023; Xue et al., 2023; Bi et al., 2023). Despite the identification of thousands of candidate regions using these approaches, only a few of these genetic changes have been shown to result in phenotypic consequences when experimentally tested in both animal and human cell culture models (Florio et al., 2015; Aldea et al., 2021; Schmidt et al., 2021; Tajima et al., 2025; Liu et al., 2025). This suggests that we may need to cast a wider net to identify important genomic sequence changes that have causal effects on human evolution.

One underexplored source of potential functional variation is tandem repeats, which are typically masked during comparative genomic analyses. Tandem repeats have a high mutation rate, allowing them to serve as a possible reservoir of evolutionary innovation (Hannan, 2018), and have previously been linked to morphological differences in dogs and sticklebacks, and to social-behavioral evolution in prairie voles (Fondon and Garner, 2004; Hammock and Young, 2005; Xie et al., 2019). However, studying tandem repeats has been limited by many technical challenges, including the difficulty of capturing entire repeat regions with short-read sequencing and of amplifying and cloning tandem repeats for functional study (Hannan, 2018). As a result, studies of tandem repeats have been mostly limited to examining short tandem repeats (STRs; 1-6 bp repeat unit) that can be covered in a single short-read sequencing read and are more amenable to routine molecular biology approaches. More recently, advances in long-read sequencing have made it possible to identify larger variable number tandem repeats (VNTRs; ≥7 bp repeat unit) that emerged during human evolution and are linked to disease risk (Sulovari et al., 2019; Course et al., 2021; Chaisson et al., 2023). However, the functional consequences of these VNTRs have rarely been tested by gain- or loss-of sequence experiments at their endogenous loci. One of the most interesting human-specific VNTRs is a large 30-mer repeat expansion in the third intron of the *CACNA1C* gene (Song et al., 2018; Sulovari et al., 2019). *CACNA1C* encodes the α subunit of the L-type voltage-gated calcium channel Ca_V_1.2. Ca_V_1.2 controls or modulates many neuronal processes, including activity-dependent transcriptional programs and dendrite morphology, and has been strongly associated with neuropsychiatric disease risk (Splawski et al., 2004; Dao et al., 2010; Jeon et al., 2010; Lee et al., 2012; Pasca et al., 2011; Krey et al., 2013; Dedic et al., 2018; Birey et al., 2022; Chen et al., 2024). The human-specific *CACNA1C* insertion is a large 3,000-30,0000+ bp VNTR with a 30 bp repeat unit that is expanded from a single 30 bp sequence found in chimpanzees. Interestingly, variation in the 30-mer sequences in modern humans is associated with risk for bipolar disorder and schizophrenia (Song et al., 2018; Moya et al., 2025), suggesting that this VNTR may play important roles in both human evolution and neuropsychiatric disease risk.

Here, we experimentally test the functional effects of this human-specific tandem repeat array, which we name TRACT for <u>t</u>andem <u>r</u>epeat <u>a</u>rray intronic to *<u>C</u>ACNA1C* with thirty-mers. We generate two complementary models: one where TRACT is inserted into the orthologous location in mice, which normally lack this repeat array; and another where TRACT is excised from human induced pluripotent stem (hiPS) cells and replaced with the single 30 bp sequence found in chimpanzees. In both the mouse and human genetic backgrounds, TRACT consistently changes the neuronal response to stimulation, likely through changes in *CACNA1C* isoform expression. A large network of transcriptional differences results from the presence or absence of TRACT in the *CACNA1C* locus, illustrating how a single VNTR may change the activity-dependent expression of neural genes in plasticity states and in the pathophysiology of neuropsychiatric diseases.

## Results

### hTRACT does not affect basal gene expression in the mouse brain

To study the effect of TRACT on human evolution, we first inserted the human TRACT sequence (hTRACT) into the orthologous location in the third intron of *Cacna1c* in mice with a mixed 129S6 and C57Bl/6J background using CRISPR/Cas9 (Fig. 1A; Materials and Methods). hTRACT was transmitted through the germline, and subsequent crosses showed that heterozygotes and homozygotes were born in expected Mendelian ratios, with normal viability and fertility. We generated two knock-in mouse lines containing distinct hTRACT alleles from different human individuals (Materials and Methods). We refer to wild-type mice, which lack hTRACT, as Mm-hTRACT^−/−^, and knock-in mice homozygous for hTRACT as Mm-hTRACT^KI/KI^.

**Figure 1.**
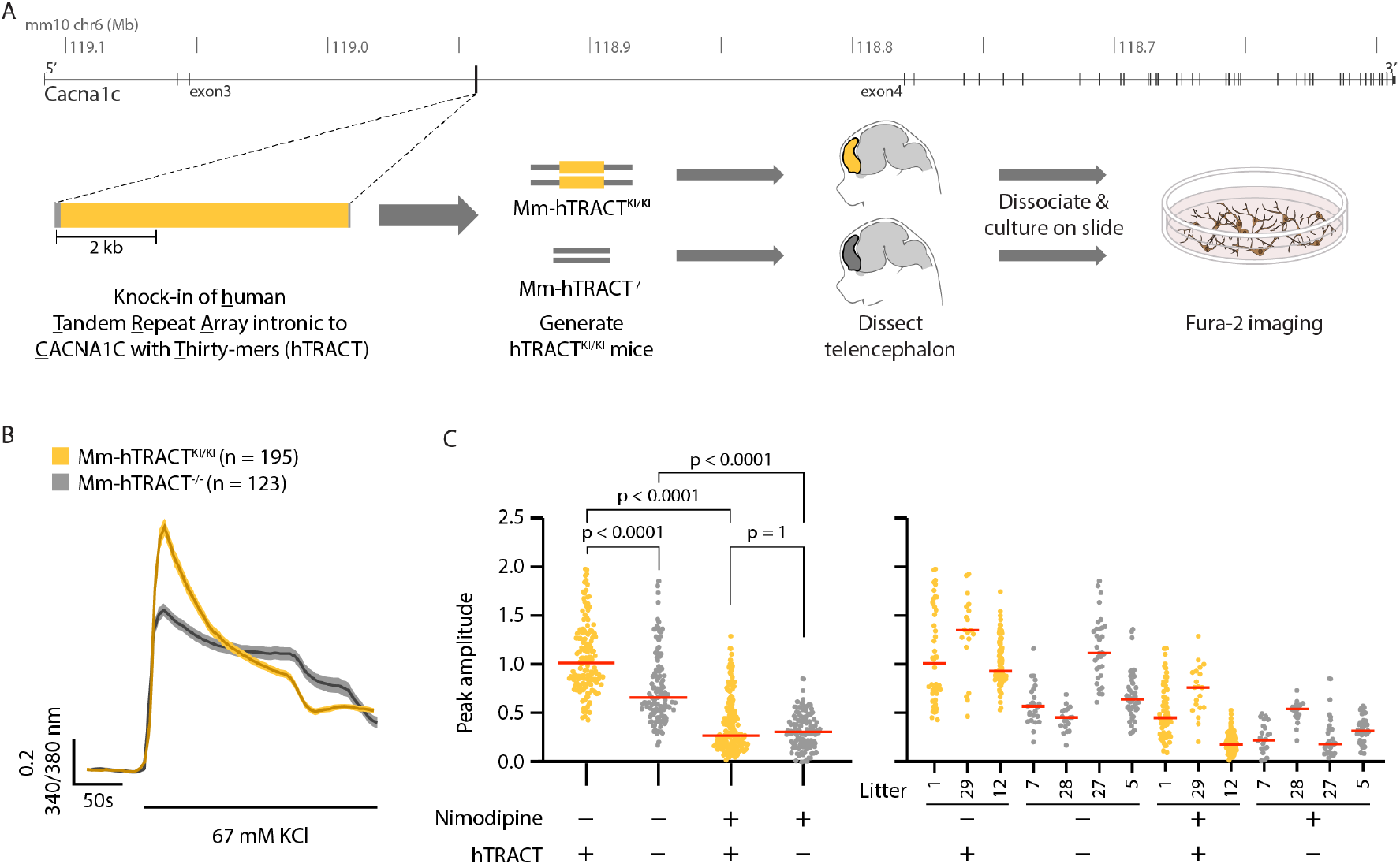
hTRACT affects calcium response to KCl stimulation in mouse neurons. (A) Schematic of the mouse model. Top: Genomic structure of mouse *Cacna1c*, with the hTRACT knock-in site in the third intron indicated by a black vertical bar. Exons are denoted as dark gray bars. Bottom: Model generation and experimental design. (B) Representative traces of intracellular calcium levels following KCl depolarization from one field of view. The y-axis represents the fluorescent intensity of the calcium reporter Fura-2. The lines and shaded areas indicate means and 95% confidence intervals, respectively. (C) Peak calcium amplitude with and without TRACT and/or nimodipine. Left, data pooled across mouse litters; right, data separated by litter. Each dot represents one cell (n = 595 cells); Kruskal–Wallis test, *p* < 0.0001. P-values of Dunn’s multiple comparison test are labeled. Red bars show median values of each group.

To determine whether hTRACT affects gene expression, we first performed bulk RNA sequencing in 14-week-old Mm-hTRACT^−/−^ and Mm-hTRACT^KI/KI^ mice. Because *Cacna1c* is widely expressed (Lonsdale et al., 2013; Baker et al., 2023), we examined the entire adult brain by subdividing it into 4 regions: the cerebral cortex and hippocampus; the midbrain, pons, and medulla; the cerebellum; and a combination of other subcortical structures including the striatum and thalamus. We sequenced 5-7 samples per region per mouse line. As expected, transcriptomes clustered by brain region along the first two principal components (Fig. S1A). To assess the effect of hTRACT, we performed differential gene expression analysis for each brain region (Table S1; Materials and Methods). Almost all of the genes that were identified as differentially expressed clustered around the location of the hTRACT insertion (Fig. S1B). This suggests that most differential genes may represent differences in gene expression between the 129S6 genetic background, where hTRACT was inserted, and the C57Bl/6J genetic background, rather than gene expression differences resulting only from hTRACT. *Cacna1c* was not identified as differentially expressed in any brain region. Outside of the locus surrounding hTRACT on chr6, only five genes were differentially expressed (Fig. S1C). Given that all five genes have adjusted p-values between 0.01 and 0.05 and that many genes are differentially expressed on chr6, it is possible that these changes also result from differences between the 129S6 and C57Bl/6J chr6 genetic backgrounds. We note that mouse lines generated on mixed backgrounds have been commonly used to study the transcriptomic effects of specific sequence changes (Yang et al., 2016), and the lack of more broadly distributed gene expression changes suggests that hTRACT does not have large effects on basal gene expression in the adult mouse brain.

To examine whether hTRACT affects brain development, we first generated two new mouse lines containing distinct hTRACT alleles in an inbred C57Bl/6J background. These lines avoid possible confounders from the mixed 129S6/C57Bl6J genetic background, and the C57Bl/6J lines were used in all subsequent experiments unless otherwise noted. We examined whether hTRACT affects gene expression in the developing brain by performing bulk RNA sequencing of the dorsal and ventral telencephalon at E12.5 and E15.5 from both mouse lines (Materials and Methods). We sequenced 6-8 samples per region per mouse line at each timepoint. As expected, the transcriptomes clustered by time point and by brain region along the first two principal components (Fig. S1D). Differential expression analysis between Mm-hTRACT^−/−^ and Mm-hTRACT^KI/KI^ mice did not identify any differentially expressed genes in the dorsal or ventral telencephalon at E12.5 or E15.5 (Fig. S1E; Table S1; Materials and Methods). Taken together, our results suggest that the insertion of hTRACT does not result in significant transcriptomic changes in the mouse brain embryonically or in adulthood.

### hTRACT perturbs calcium signaling following stimulation in mouse neurons

Given the key role of Ca_V_1.2 in mediating the intracellular response to neuronal activity (Simms and Zamponi, 2014), we wondered whether hTRACT might affect responses of cultured neurons to chemical stimulation by potassium chloride (KCl), which depolarizes membranes, leading to activation of calcium channels, and is a commonly used model to study activity-dependent processes (Sheng and Greenberg, 1990; Tao et al., 1998). We examined intracellular calcium signaling following depolarization in neurons that had been dissociated from the telencephalon of Mm-hTRACT^−/−^ and Mm-hTRACT^KI/KI^ mice at E15.5 and grown in culture for 7-10 days (Materials and Methods). Dissociated cultures were infected with an AAV encoding the eYFP reporter under the control of the human Synapsin-1 (hSYN1) promoter to label neurons. We used the ratiometric Fura-2 calcium indicator and depolarized neurons with 67 mM KCl (Fig. 1B). Strikingly, Mm-hTRACT^KI/KI^ neurons had a significantly higher amplitude of normalized Fura-2 signal, indicative of perturbed calcium signaling response to depolarization, compared to Mm-hTRACT^−/−^ neurons upon depolarization (*p* < 0.0001; Fig. 1B, Fig. S2A).

We then verified whether this change in calcium signaling is mediated by L-type voltage-gated calcium channels, including Ca_V_1.2. Exposing both Mm-hTRACT^−/−^ and Mm-hTRACT^KI/KI^ neurons to nimodipine, which inhibits L-type voltage-gated calcium channels, significantly decreased the Fura-2 signal amplitude upon depolarization (*p* < 0.0001; Fig. 1C, Fig. S2B). There was no significant difference in Fura-2 signal amplitude between Mm-hTRACT^KI/KI^ and Mm-hTRACT^−/−^ neurons in the presence of nimodipine (*p* = 1). This suggests that hTRACT affects intracellular calcium amplitude upon neuronal depolarization through L-type voltage-gated calcium channels such as Ca_V_1.2.

### TRACT alters depolarization-induced calcium signaling in human cortical neurons

Because mice and humans are separated by 90 million years of evolution (Hedges et al., 2006), we also tested the effect of TRACT on the neuronal response to stimulation in a human genetic background. Using CRISPR/Cas9 editing, we generated isogenic hiPS cell lines where TRACT was excised from the endogenous human *CACNA1C* locus and replaced with the single 30 bp sequence found in chimpanzees (Hs-TRACT^−/−^) on both alleles (Fig. 2A; Materials and Methods). Using the hiPS cell line 6032-4, we generated three Hs-TRACT^−/−^ lines, as well as four unedited control lines (Hs-TRACT^+/+^) that went through the same steps of the CRISPR/Cas9 protocol. We then differentiated matched isogenic hiPS cell lines with and without TRACT into human cortical organoids as we have previously described (Pasca et al., 2015; Yoon et al., 2019).

**Figure 2.**
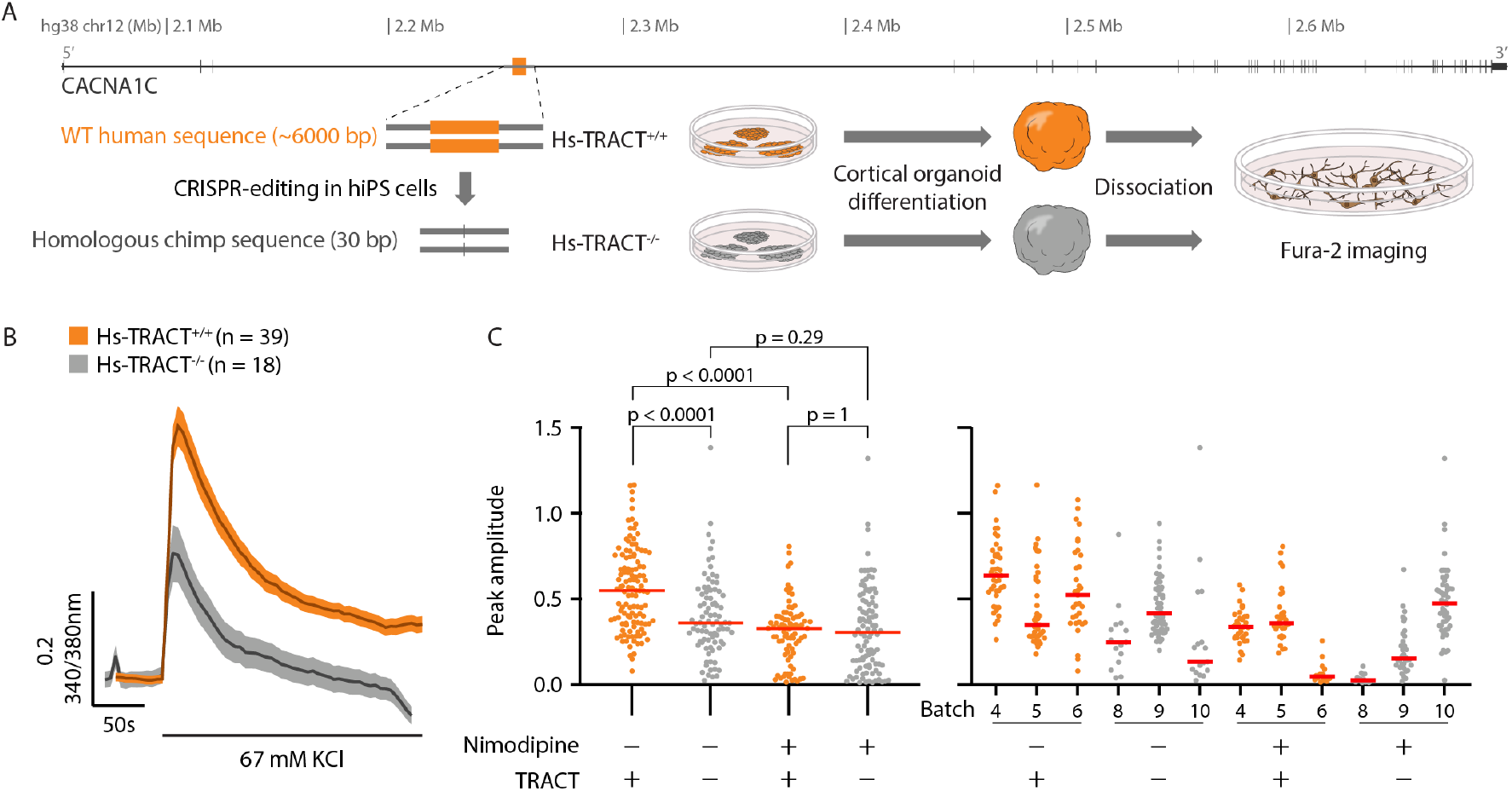
TRACT affects calcium response to KCl stimulation in hiPS cell-derived neurons. (A) Schematic of the human cortical organoid model. Top: Genomic structure of human *CACNA1C*, with TRACT in the third intron highlighted as an orange box. Exons are denoted as dark gray bars. Bottom: Model generation and experimental design. (B) Representative traces of intracellular calcium levels following KCl depolarization from one field of view. The y-axis represents the fluorescent intensity of the calcium reporter Fura-2. The lines and shaded areas indicate means and 95% confidence intervals, respectively. (C) Peak calcium amplitude with and without TRACT and/or nimodipine. Left, data pooled across biological replicates; right, data separated by replicate. Each dot represents one cell (n = 384 cells); Kruskal–Wallis test, *p* < 0.0001. P-values of Dunn’s multiple comparison test are labeled. Red bars show median values of each group.

Human cortical organoids (hCO) were dissociated and plated at day 188-254 (D188-254) of differentiation and grown in plated 2D culture for 20-30 days prior to imaging. Neurons were labeled with an AAV expressing eYFP under the hSYN1 promoter (Materials and Methods). Consistent with the change in intracellular Fura-2 amplitude between Mm-hTRACT^KI/KI^ and Mm-hTRACT^−/−^ neurons, Hs-TRACT^+/+^ neurons had a significantly higher calcium signal than Hs-TRACT^−/−^ neurons upon depolarization with 67 mM KCl (*p* < 0.0001; Fig. 2B). This increase in intracellular calcium signaling was not observed when human neurons were exposed to nimodipine (*p* = 1; Fig. 2C). These results suggest that the presence of TRACT increases intracellular calcium levels after chemical depolarization in both mouse and human cortical neurons.

### TRACT causes correlated changes in activity-dependent gene expression in both mouse and human neurons

Calcium influx following depolarization induces an activity-dependent transcriptional response that drives downstream cellular and circuit-level changes (Yap and Greenberg, 2018). This includes changes in the expression of immediate early genes within 15 minutes (min) that peaks at ~1 hour (h). Immediate early genes then drive a second transcriptional wave of late response genes that peaks ~6 h after calcium influx (Yap and Greenberg, 2018). To examine the effect of TRACT on both immediate early and late response gene programs, neurons dissociated from the mouse telencephalon at E15.5 or hCO at D188-D254 were grown in culture, exposed to 67 mM KCl for 1 h or 6 h, collected, and processed for bulk RNA sequencing (Fig. 3A, Materials and Methods). Control neurons from the same mice or hCO that had not been exposed to KCl were collected and processed concurrently. We additionally generated two Hs-TRACT^−/−^ lines and two control Hs-TRACT^+/+^ lines derived from the hiPS cell line H20961 as a second set of isogenic lines to use in these experiments.

**Figure 3.**
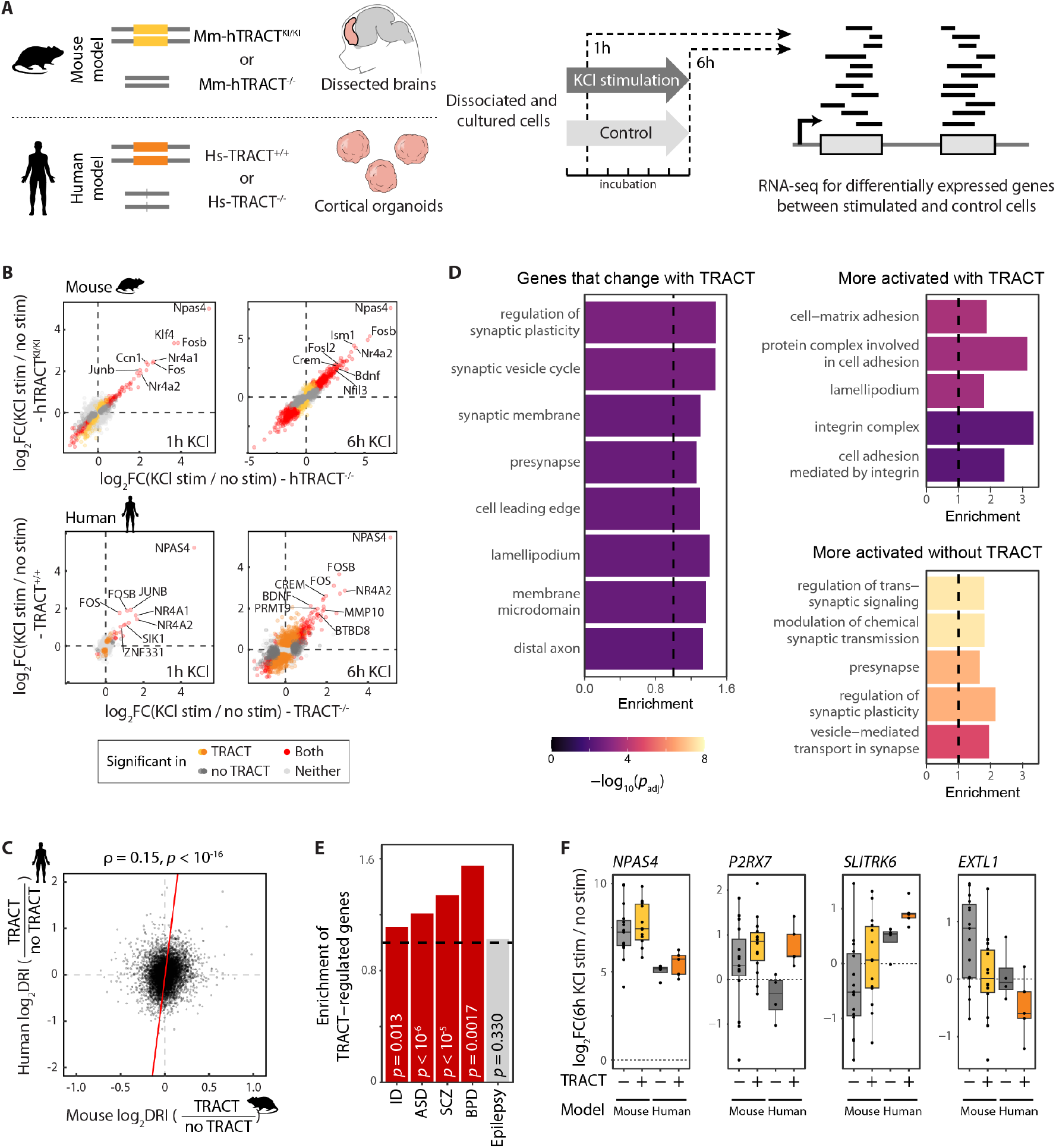
TRACT causes concordant changes in activity-dependent transcription after KCl stimulation in mouse and human neurons. (A) Schematic of the experimental design. (B) Transcriptional response to 1 h (left) or 6 h (right) of KCl stimulation (stim) in cells with TRACT (y-axis) versus cells without TRACT (x-axis). (C) Differential regulation indices (DRI) for mouse and human are significantly correlated for activity-dependent genes after 6 h of KCl stimulation (Materials and Methods). (D) Gene ontology terms for activity-dependent genes that have the same direction of effect with TRACT in both mouse and human models. Left: All genes. Right: Genes that have a stronger or weaker activity-dependent transcriptional response with TRACT. (E) Genes with the same direction of effect due to TRACT in both mouse and human models are enriched for intellectual disability (ID) and autism spectrum disorder (ASD) genes (Abrahams et al., 2013; Kochinke et al., 2016) and are enriched near schizophrenia (SCZ) and bipolar disorder (BPD) GWAS loci (Mullins et al., 2021; Trubetskoy et al., 2022), but are not enriched for epilepsy genes (Leblond et al., 2021). (F) Examples of genes where TRACT has concordant effects on activity-dependent expression after 6 h KCl stimulation between mouse and human neurons.

At the early time point (1 h), canonical immediate early genes, including *Fos, Npas4*, and *Nr4a1* (Yap and Greenberg, 2018), were strongly up-regulated in both mouse and human neurons (Fig. 3B; Table S2). At the later time point (6 h KCl), we observed thousands of activity-dependent genes in both mouse and human neurons compared to 1 h KCl (Fig. 3B; Table S2), consistent with prior studies (Ataman et al., 2016).

To examine whether TRACT affects activity-dependent gene expression, we calculated the differential regulation index (DRI) between neurons with and without TRACT. DRI has been previously used to calculate the change in activitydependent transcription between human and mouse neurons, DRI(Hs/Mm), by dividing the fold change after KCl treatment in human neurons by the fold change after KCl treatment in mouse neurons (Qiu et al., 2016). Analogously, we calculated DRI(TRACT / no TRACT) for both mouse (Mm-hTRACT^KI/KI^ / Mm-hTRACT^−/−^) and human (Hs-TRACT^+/+^ / Hs-TRACT^−/−^) neurons and examined whether DRI(TRACT / no TRACT) was correlated between the two models, which would indicate that TRACT leads to consistent changes in activity-dependent gene expression across genetic backgrounds.

Although DRI(TRACT / no TRACT) was not correlated between mouse and human neurons at the 1 h KCl time point (Fig. S3A), we observed a modest but highly significant correlation at the 6 h KCl time point (Spearman’s ρ = 0.15, *p* < 10^−16^) (Fig. 3C). Compared to all expressed genes, genes with concordant changes in activity-dependent expression (same direction of effect in both mouse and human neurons) were enriched for the gene ontology terms regulation of synaptic plasticity, the synaptic membrane, and cell leading edge (Fig. 3D, left; Table S2). Additionally, genes that were **more** activity-regulated in the presence of TRACT were enriched for ontology terms involved in cell adhesion, particularly by the integrin complex (Fig. 3D, top right; Table S2), which is known to modulate calcium signaling and growth cone extension (Wu and Reddy, 2012). In contrast, genes that were **less** activity-regulated in the presence of TRACT were enriched for ontology terms related to synaptic transmission (Fig. 3D, bottom right; Table S2).

Non-coding intronic variants that map near or in TRACT itself have been linked to human neuropsychiatric conditions, including bipolar disorder and schizophrenia (Ferreira et al., 2008; Ripke et al., 2011; Song et al., 2018; Moya et al., 2025). Having identified a large set of genes whose activity-dependent expression in neurons differs when TRACT is present in the genome, we asked whether these TRACT-dependent genes are also disease-associated. We find that the set of 4062 genes whose expression is consistently affected by TRACT are also significantly enriched near GWAS loci previously associated with bipolar disorder (fold enrichment = 1.55; adjusted *p* = 0.0017) and schizophrenia (fold enrichment = 1.34; adjusted *p* < 10^−5^) (Fig. 3E; Materials and Methods). Weaker enrichment is seen with genes previously associated with autism spectrum disorder (fold enrichment = 1.21; adjusted *p* < 10^−6^) and intellectual disability (fold enrichment = 1.11; adjusted *p* = 0.013). In contrast, no significant enrichment is seen for genes previously implicated in epilepsy (fold enrichment = 1.02; *p* = 0.330). Thus, multiple common neurological conditions are enriched for TRACT-dependent genes, with the most pronounced enrichments seen for common neuropsychiatric diseases like bipolar disorder and schizophrenia that are associated not only with genetic variation in and near TRACT itself, but also with genetic variation in many other genes whose activity-dependent expression is now known to be modulated by TRACT.

We highlight a few genes where the presence of TRACT affects the degree of activity regulation (Fig. 3F): *NPAS4, P2RX7*, and *SLITRK6*, where TRACT increases expression, and *EXTL1*, where TRACT decreases expression. *NPAS4* is a canonical immediate early gene that acts as a transcription factor to activate the expression of late response genes (Yap and Greenberg, 2018). Although *NPAS4* expression does not consistently change with TRACT at 1 h KCl (Fig. S3B), increased *NPAS4* expression at 6 h KCl could indicate a prolonged period of *NPAS4* expression and consequently, prolonged activation of its transcriptional targets. In contrast, *P2RX7, SLITRK6*, and *EXTL1* are not activity-dependent at 1 h KCl, act as late response genes at 6 h KCl, and may affect synapse formation. The neuropsychiatric disease gene *P2RX7* is a purinergic receptor that affects axonal outgrowth and dendritic branching (Miras-Portugal et al., 2017; Mut-Arbona et al., 2023), *SLITRK6* is a synaptic transmembrane protein implicated in neurite outgrowth (Katayama et al., 2009; Proenca et al., 2011), and *EXTL1* encodes a glycosyltransferase essential for polymerization of heparan sulfate (Kim et al., 2001), a glycan critical for synapse formation (Zhang et al., 2018). We note that differences in the degree of activity-dependent expression for *SLITRK6* and *EXTL1* between mouse and human neurons could be due to differences in the mouse and human genetic backgrounds or differences in developmental timing and cell type identity specific to experimental differences between the mouse and human models. Nevertheless, our results suggest that TRACT drives global changes to the activity-dependent transcriptional program across both mouse and human models, affecting genes involved in neurite outgrowth and synaptic signaling.

### TRACT recapitulates known differences in activity-dependent gene expression between mouse and human neurons

Unlike immediate early genes, which are largely consistent across neuronal maturation and different neuronal subtypes, late response genes can vary with developmental timing, subtype identity, and species (Hrvatin et al., 2018; Traunmüller et al., 2025; Carter et al., 2025). To identify a core set of late response genes that are consistently affected by TRACT, we first identified late response genes where the degree of activity dependence consistently changes between wild-type human and mouse neurons, i.e. where DRI(Hs/Mm) have the same sign. We examined activity-dependent gene expression between the wild-type human and mouse neurons from this study (Hs-TRACT^+/+^ / Mm-hTRACT^−/−^), as well as from two prior studies that examined differences in activity-dependent transcription between human and mouse neurons (Ataman et al., 2016; Qiu et al., 2016). We reasoned that focusing on late response genes with consistent changes between human and mouse neurons would identify evolutionarily relevant late response genes that likely act across a range of developmental windows and neuronal subtypes. Of the 6,208 late response genes in this study that were also examined in the two other studies, DRI(Hs/Mm) was consistent across all three datasets for 2,448 late response genes with the remainder divided between those with consistent effects in two out of the three datasets (Fig. S3C; Table S2).

Because TRACT is a human-specific insertion, we focused on late response genes where DRI(TRACT / no TRACT) has the same sign in both mouse and human neurons and where DRI(TRACT / no TRACT) also matches the sign of DRI(Hs/Mm) across all three datasets. In Fig. 4A, we highlight the 36 late response genes with the largest changes in DRI(TRACT / no TRACT). These include genes that have an increased activity-dependent response in both human and mouse neurons with TRACT, such as *CREB3L2*, a transcriptional activator that can promote axon outgrowth (McCurdy et al., 2019) (Fig. 4B), and *GABRG3*, a GABA receptor subunit associated with neurodevelopmental disorders (Wang et al., 2018; Frohlich et al., 2019). Other genes, such as the oncogene *WDR54* (Yuan et al., 2019) and the canonical immediate early gene *EGR1* (Yap and Greenberg, 2018), have decreased activity-dependent responses in both human and mouse neurons with TRACT (Fig. 4B). We also observe many genes with the same direction of change but different baselines between mouse and human neurons, including the tumor suppressors *PDCD7* (Peng et al., 2018) and *GADD45G* (Ying et al., 2005) (Fig. 4B). *PDCD7* is negatively activity-regulated in wild-type mouse neurons but is weakly, if at all, activity-regulated in wild-type human neurons; neurons containing TRACT have weakened activity-regulation relative to neurons of the same species that do not contain TRACT, indicating that TRACT contributes to a decreased response to activity (Fig. 4B). Similarly, *GADD45G* is positively activity-regulated in wild-type mouse neurons with weakened activity regulation in control human neurons, and TRACT contributes to this weakened response to activity (Fig. 4B). Intriguingly, prior work suggests that *GADD45G* is also regulated by a *cis*-acting human-specific deletion, although the activity-dependence of this non-coding element has not been assessed (McLean et al., 2011). Differences in activity regulation between human and mouse neurons for specific genes like *PDCD7* and *GADD45G* are likely due to a composite of *trans* effects, through elements like TRACT, and *cis* effects, such as the human-specific deletion near *GADD45G*.

**Figure 4.**
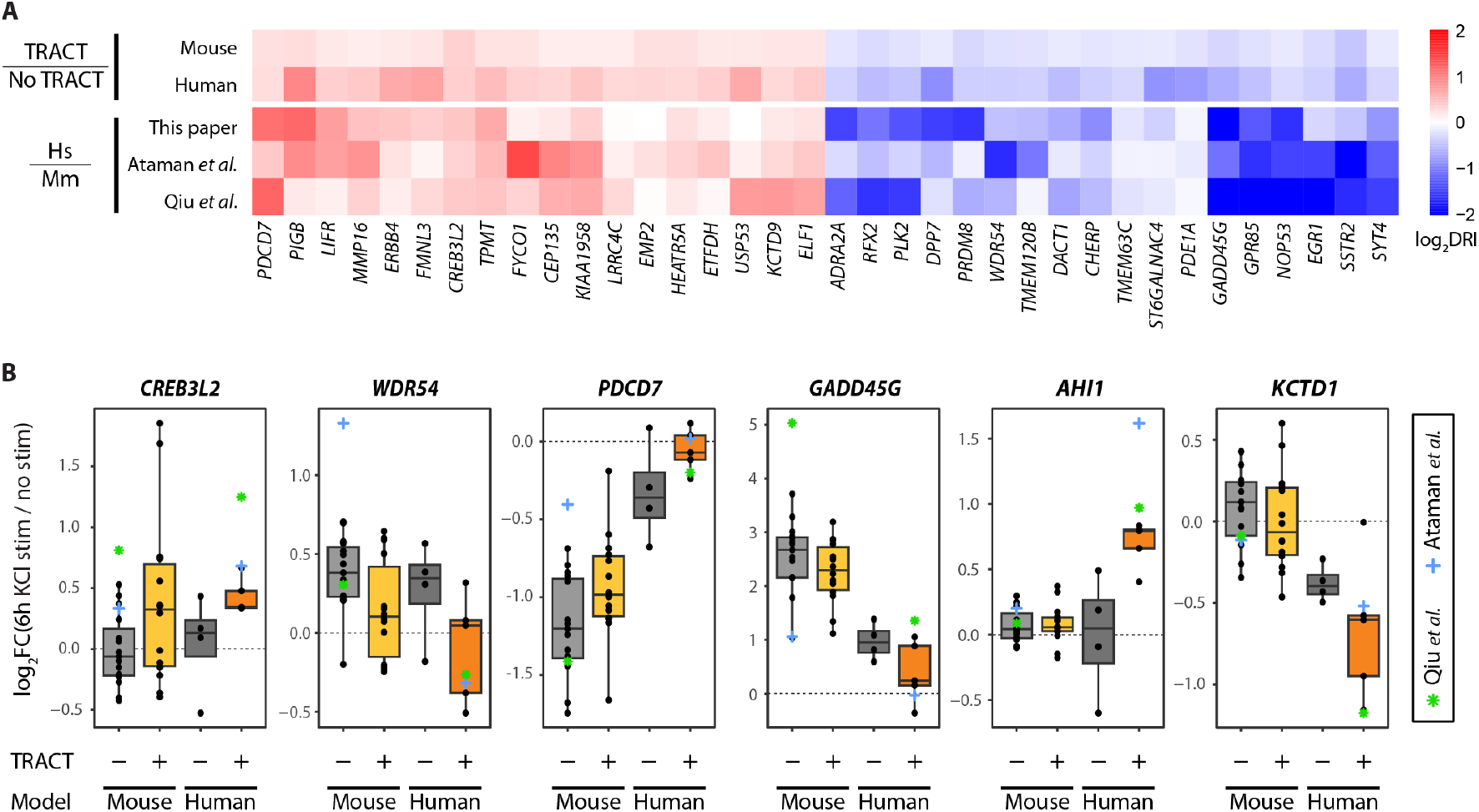
TRACT contributes to known differences in activity-dependent transcription between mouse and human neurons. (A) DRI heatmap of representative genes with concordant changes between TRACT vs. no TRACT comparisons in mouse and human neurons after 6 h KCl stimulation (top two rows) and wild-type human vs. wild-type mouse neurons in this study and two previous publications after 4-6 h KCl stimulation (bottom three rows). (B) Fold changes of selected genes from (A). Fold changes reported in previous publications are labeled as green asterisks for (Qiu et al., 2016) and blue crosses for (Ataman et al., 2016).

Although DRI(TRACT / no TRACT) is significantly correlated between human and mouse neurons, the magnitude of DRI(TRACT / no TRACT) is much larger in human compared to mouse neurons (slope = 15.2). In addition, many of the late response genes affected by TRACT, including *PDCD7* and *GADD45G*, have consistent differences in activity-dependent expression between wild-type mouse and human neurons across our dataset and prior datasets (Fig. 4B). This suggests that differences between mouse and human neurons lacking TRACT, or between mouse and human neurons with TRACT, cannot be completely explained by experimental differences in developmental timing or cell type composition but likely reflect species-specific differences in activity-dependent transcription. Indeed, we observe that *OSTN*, a secreted factor, and *ZNF331*, a transcriptional repressor with no mouse ortholog, which have both been previously found to be activity-regulated in human but not in rodent neurons (Ataman et al., 2016), are also activity-regulated in our human but not in mouse neurons regardless of TRACT status (Table S2).

TRACT may also have effects specific to the human but not the mouse genetic background. For example, we find that *AHI1*, a ciliary transition zone gene (Hsiao et al., 2021) that has been previously identified as activity-dependent in human but not rodent neurons (Ataman et al., 2016), is activity-dependent with, but not without, TRACT in human neurons, suggesting that TRACT is necessary for its human-specific expression pattern. However, TRACT alone is not sufficient for the activity-dependent expression of *AHI1*, as *AHI1* is not activity-dependent in mouse neurons with or without TRACT (Fig. 4B). Similarly, *KCTD1*, a transcriptional repressor (Ding et al., 2008), is not activity-regulated in wild-type mouse neurons but is negatively activity-regulated in wild-type human neurons (Ataman et al., 2016; Qiu et al., 2016). This negative regulation is attenuated but not abolished in human neurons that lack TRACT, suggesting that TRACT contributes to the activity-dependent regulation of *KCTD1* observed specifically in humans (Fig. 4B). Taken together, our results suggest that TRACT drives specific changes in activity-dependent transcription in both mouse and human neurons, with stronger effects in human neurons due to differences between the mouse and human genetic backgrounds.

### TRACT shifts the relative abundance of specific CACNA1C isoforms

How does TRACT, a non-coding tandem repeat found in the third intron of *CACNA1C*, lead to changes in calcium signaling and activity-dependent transcription? RNA sequencing and allele-specific expression studies did not detect an effect of TRACT on total *CACNA1C* RNA expression levels, either in mouse brain regions collected at multiple embryonic and postnatal stages, or in hCO collected at several time points (Fig. S1, Fig. S4, Fig. S5; Table S1, S3; Materials and Methods). However, the *CACNA1C* gene is known to have hundreds of different splicing isoforms (Clark et al., 2020), and these isoforms perform varied functions in development and in disease (Splawski et al., 2005; Panagiotakos et al., 2019; Harrison et al., 2022).

To further examine *CACNA1C* isoform expression, we performed long-read amplicon sequencing of *CACNA1C* in mouse telencephalon and hCO (Clark et al., 2020; Hall et al., 2021) (Table S4; Materials and Methods). In both models, we identified known and novel *CACNA1C* exons and splice junctions when compared to those currently listed in the NCBI refSeq database or identified in previous studies (Clark et al., 2020) (Fig. S6). While the fraction of reads with novel human exons was <0.02%, the fraction of reads with novel mouse exons was 20.05% (Fig. S6A). The higher abundance of newly described exons in mice likely reflects fewer past research efforts to profile *CACNA1C* isoforms in mice compared to humans.

To identify potential effects of TRACT on *CACNA1C* isoform expression, we identified orthologous exons between the mouse and human annotations, determined the fraction of transcripts that contain each exon, and compared relative abundances between samples with and without TRACT (Materials and Methods). While most orthologous exons are found in nearly all transcripts and show little variation, exon 31 and exon 32 are alternative exons found in 34% and 63% of transcripts respectively (Fig. 5A, Fig. S6B). We found that exon 31 trended to higher abundance in samples with TRACT compared to samples without TRACT (24% increase, *p* = 0.127) and conversely that exon 32 was 8% less abundant (*p* = 0.029; Fig. 5B).

**Figure 5.**
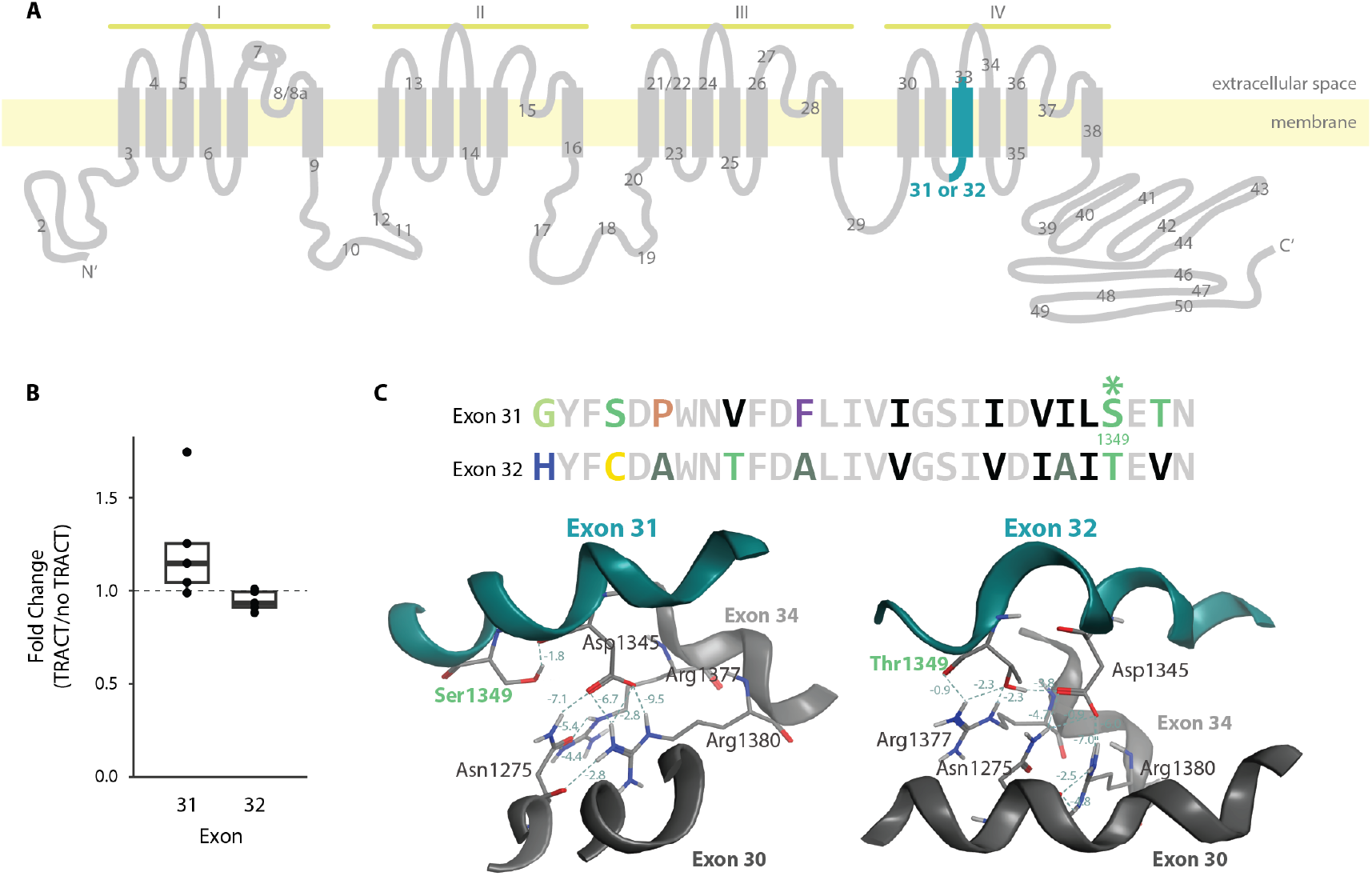
TRACT may affect *CACNA1C* exon usage. (A) Schematic of the Ca_v_1.2α_1c_ subunit, the protein product of *CACNA1C* (not to scale). Numbers along the peptide mark the start of the exon. The segment corresponding to alternative exons 31/32 is highlighted in teal. Roman numerals denote transmembrane domains. (B) Fold changes for exons 31 and 32 between samples with and without TRACT. Dashed line denotes no change. (C) Top: Amino acid sequences of exons 31 and 32 with differential residues highlighted and colored according to chemical properties. Bottom: Predicted structural differences between Ca_v_1.2α_1c_ with Ser1349 (exon 31) and Thr1349 (exon 32). Predicted hydrogen bonds and their strengths are indicated by light blue dashed lines and numbers.

To gauge the possible functional consequences of alternative exon 31/exon32 usage, we examined differences between the exon 31 and exon 32 protein sequences. The Ca_v_1.2α_1c_ subunit, the protein product of *CACNA1C*, includes 4 major domains (I – IV), each with 6 transmembrane domains (S1 – S6) (Tang et al., 2004). In domain IV, exons 31/32 encode the S3 transmembrane domain and form a voltage-sensing unit with the S4 transmembrane domain (encoded by exon 34). Modeling differences between exons 31 and 32 using the solved structure of Ca_v_1.2 (Chen et al., 2023) identified three residue differences that are predicted to alter hydrogen bonds with neighboring amino acids and one residue change that is predicted to alter a pi-cation bond (Fig. S6D). In particular, a change from Thr1349 in exon 32 to Ser1349 in exon 31 is predicted to strengthen hydrogen bonding between the aspartic acid, Asp1345, in the S3 transmembrane domain and the arginines, Arg1377 and Arg1380, in the S4 transmembrane domain (Fig. 5C). Strikingly, loss of interactions between Asp1345, Arg1377, and Arg1380 have been previously shown to lower channel open probability and current density (Costé de Bagneaux et al., 2018). This raises the possibility that strengthened interactions between these residues in humans may increase current density and open probability, consistent with the changes in calcium amplitude we observed in neurons with TRACT.

## Discussion

In this study, we used reciprocal genetic modifications to study the effect of TRACT, a human-specific VNTR intronic to *CACNA1C*, in both mice and human cortical organoids. We find that presence of TRACT increases the amplitude of calcium signaling following chemical depolarization in both mouse and human neurons and leads to large-scale concordant changes in activity-regulated transcription. Strikingly, genes for which TRACT alters the transcriptional response to activity are enriched for those that are involved in synaptic functions, and are also enriched near genes implicated in multiple human neuropsychiatric diseases.

What might be the consequences of TRACT changes to neuronal stimulation? Humans have a prolonged period of neuronal maturation and synapse refinement that extends into postnatal life. This elongated developmental period is critical for proper circuit formation and cognition, and appears to be a species-specific, cell-intrinsic feature that is maintained in neural transplants and cell culture (Gaspard et al., 2008; Petanjek et al., 2011; Espuny-Camacho et al., 2013; Otani et al., 2016; Sloan et al., 2017; Trevino et al., 2020; Gordon et al., 2021; Revah et al., 2022). We find that the large network of genes that are more activity-regulated when TRACT is present at the *CACNA1C* locus are enriched for cell adhesion genes, like integrins, that promote axon extension and neurite outgrowth. In contrast, the set of genes that are less activity-regulated when TRACT is present at the *CACNA1C* locus are enriched for processes involved in synaptic transmission. The overall shift toward genes associated with neurite outgrowth and away from genes involved in trans-synaptic signaling suggests that TRACT might be one of the genomic mechanisms contributing to a prolonged developmental window for neuronal maturation in humans.

Although previous studies have shown that TRACT sequences have enhancer activity in cultured cells (Song et al., 2018), we find that presence or absence of TRACT does not change total *CACNA1C* RNA levels but instead increases the proportion of *CACNA1C* isoforms containing exon 31 (versus the alternative exon 32). Isoforms containing exon 31 are more highly expressed during development, whereas isoforms containing exon 32 are more highly expressed in adulthood (Diebold et al., 1992; Clark et al., 2020; Mazin et al., 2021). Thus, an increase in isoforms containing exon 31 would be consistent with TRACT elongating the period of neuronal maturation. Mechanistically, exon 31/32 and exon 34 encode two transmembrane domains that interact to form a voltage-sensing unit in Ca_v_1.2, and we find that amino acid differences between exon 31 and 32 are predicted to change hydrogen bonding between key residues that have been previously found to affect channel properties (Costé de Bagneaux et al., 2018). Although this provides a possible explanation for how TRACT leads to differences in the response to neuronal stimulation, additional experimentation will be needed to test whether increased expression of isoforms containing exon 31 can phenocopy the effects seen in mouse and human TRACT models.

TRACT may also operate via alternate mechanisms. For example, previous studies have shown that both genic and intergenic tandem repeat expansions can be transcribed and form RNA foci that sequester RNA binding proteins and perturb cellular function and viability (Zhang and Ashizawa, 2017). TRACT also has potential non-ATG open reading frames in both directions (Song et al., 2018), and other tandem repeats are known to be translated even in the absence of conventional ATG start codons (Zu et al., 2011; Cleary and Ranum, 2013; Banez-Coronel et al., 2015).

Our results demonstrate that presence or absence of a single VNTR that has evolved in the human genome can lead to striking and reproducible changes in both calcium signaling and activity-dependent transcriptional responses to neural stimulation. Despite great interest in genomic changes that underlie human evolution, very few specific DNA alterations have been confirmed to produce significant phenotypic effects when recreated at the corresponding locus in both animal and human systems (Liu et al., 2025). Our study highlights the value of modeling genomic changes by both recapitulating the sequence change in mice, and reverting the sequence change in human cells. Although the effect size of TRACT on activity-dependent transcription was weaker in the mouse genetic background, a mouse model is a fully intact animal system that provides a platform for future behavioral experiments. In contrast, the human cortical organoid model matches the genetic background on which TRACT evolved, but grows without the long-range neural inputs and structural support provided by blood vessels, the skull, and other organs in animal models. The concordant changes seen when either adding TRACT to the mouse *Cacna1c* locus or removing TRACT from the human *CACNA1C* locus, suggest that the complementary mouse and human models can provide a powerful basis for studying the phenotypic effects of human-specific sequence changes.

Further, our work demonstrates that VNTRs can have profound functional effects on neural traits. Although it has been speculated that VNTRs play important roles in evolution and in disease due to their high mutation rate, accurately identifying VNTRs has been difficult prior to the advent of long-read sequencing technologies (Hannan, 2018). Recent examination of human and non-human primates with long-read sequencing has 9 now catalogued >1000 VNTRs that have expanded in the human lineage (Sulovari et al., 2019), but functional tests of these VNTRs have been limited. Our finding that TRACT affects the neuronal response to stimulation strongly motivates future efforts to study VNTRs for potential roles in the evolution of human-specific traits.

In addition to its role in human evolution, TRACT variants within modern human populations are linked to some of the most highly significant and widely replicated GWAS markers for bipolar disorder and schizophrenia (Ferreira et al., 2008; Ripke et al., 2011; Song et al., 2018; Mullins et al., 2021; Trubetskoy et al., 2022; Moya et al., 2025). Our studies provide compelling new evidence that TRACT is a key functional element in the human *CACNA1C* locus. Future work modeling different TRACT variants associated with either protection or risk from bipolar disorder and schizophrenia in both mouse and human models can now be carried out to assess how human sequence variation in TRACT may affect neuropsychiatric disease risk. Increasing evidence supports the hypothesis that human-specific variants that contribute to the evolution of human-specific traits may also preferentially contribute to diseases that affect those traits (Srinivasan et al., 2016; Shin et al., 2024). Further investigation of TRACT and its genetic variability within modern humans may provide valuable insight into both the evolution of altered neural functions in the human lineage, and human susceptibility to neuropsychiatric diseases.

## Acknowledgements

We thank members of the Kingsley and Pasca labs for helpful discussions. Caiying Guo at Janelia Research Campus assisted with the generation of the mouse models, and Kyomi Igarashi assisted with the pyrosequencing experiments.

## Funding

This work was supported by the National Science Foundation Graduate Research Fellowship (JHTS), Stanford Graduate Fellowship (JHTS), Stanford CEHG Fellowship (JHTS), and UKRI BBSRC (BB/CCG1720/1, BS/E/T/000PR981, BBS/E/ER/230001A, BBS/E/ER/230001C) (WH). DMK is an Investigator of the Howard Hughes Medical Institute.

## Author contributions

**Janet H.T. Song**: Conceptualization, Methodology, Investigation, Visualization, Writing - Original Draft, Writing - Review & Editing. **Fikri Birey**: Conceptualization, Methodology, Investigation, Visualization, Writing - Original Draft, Writing - Review & Editing. **Tzu-Chiao Hung**: Methodology, Investigation, Visualization, Writing - Original Draft, Writing - Review & Editing. **Nicola A.L. Hall**: Methodology, Investigation, Writing - Original Draft, Writing - Review & Editing. **Catherine A. Guenther**: Investigation, Writing - Review & Editing. **Xiaoyu Chen**: Investigation. **Ibrahim F. Alkuraya**: Investigation, Visualization, Writing - Original Draft, Writing - Review & Editing. **Elizabeth M. Tunbridge**: Methodology, Writing - Review & Editing. **Wilfried Haerty**: Methodology, Investigation, Writing - Original Draft, Writing - Review & Editing. **Sergiu P. Pasca**: Conceptualization, Methodology, Writing - Original Draft, Writing - Review & Editing. **David M. Kingsley**: Conceptualization, Methodology, Writing - Original Draft, Writing - Review & Editing.

## Competing interests

Elizabeth Tunbridge is a full time employee of Boehringer Ingelheim. Prior to her move into industry, she was in receipt of unrestricted research grants from J&J Innovation, and from Boehringer Ingelheim and Biogen, via the Psychiatry Consortium of the Medicines Discovery Catapult. She also provided consultancy to Boehringer Ingelheim and ONO Pharma. She reports no conflict of interest with the current manuscript. The remaining authors declare that they have no competing interests.

## Data and Materials Availability

Sequencing data is available on GEO at GSE286408. Cell lines and mouse models are subject to MTAs and available upon request. All other data are available in the main text or the supplementary materials.

