## Supplemental Materials for "Human-specific tandem repeat in *CACNA1C* modulates responses to neuronal stimulation"

### Materials and Methods

*Generation of knock-in mouse lines*

We separately cloned both a human repeat allele with 192 30-mer repeats (GenBank accession number MH645926.1) and a human repeat allele with 190 30-mer repeats (GenBank accession number MH645944.1) into the pJAZZ linear vector (Lucigen) as previously described (Song et al., 2018). Because the LA Taq DNA Polymerase (ClonTech) required to amplify the human tandem repeat is a Taq polymerase that lacks proofreading ability, we screened a total of 657 clones (407 for the 192-copy tandem repeat and 250 for the 190-copy tandem repeat) to identify clones where the inserted sequence exactly matched the expected sequence. These clones were first screened via restriction digest to check the size of the insertion and via Sanger sequencing to check the repeat sequence on the 5' and 3' end of the insertion. 115 clones (60 for the 192-copy tandem repeat and 55 for the 190-copy tandem repeat) passed initial filtering and were sequenced via long-read PacBio sequencing and analyzed as described previously (Song et al., 2018). Out of these clones, one clone each exactly matched the sequence of the two original human alleles.

The tandem repeats in these verified clones were then moved using restriction digestion with AscI and NotI into a vector containing mouse homology arms from the homologous mouse locus in the third intron of *CACNA1C* and a neomycin resistance gene surrounded by FRT sites (Fig. S7). The sequences of the tandem repeats were again confirmed via long-read PacBio sequencing. Mice containing the human tandem repeat were then generated by homologous recombination in mouse embryonic stem cells. The insertion was transmitted through the germline, and bred to mice expressing Flp recombinase (Jackson Laboratory, cat #009086) to excise the neomycin resistance cassette. The mice expressing Flp recombinase had been back-crossed to C57Bl/6J for at least 13 generations. In addition to the tandem repeat, 166 bp of additional sequence was inserted into the homologous mouse locus (Fig. S7). The sequences of the tandem repeat insertions were verified by long-read PacBio sequencing. Genotyping primers are found in Table S5.

Knock-in mice containing the human tandem repeats were generated twice, once on a mixed 129S6 and C57Bl/6 background and once on an inbred C57Bl/6J background. The tandem repeat inserted on the 129S6 allele when generated in the mixed background. All lines were back-crossed to C57Bl/6J. Wild-type mouse lines were referred to as Mm-hTRACT$^{-/-}$, and knock-in mouse lines were referred to as Mm-hTRACT$^{KI/KI}$. Mouse work was carried out in accordance with protocols approved by the Stanford University Institutional Animal Care and Use Committee (APLAC-10665).

*Culture and neural differentiation of hiPS cell lines*

The hiPS cell lines used in this study were validated using standardized methods as previously described (Yoon et al., 2019). hiPS cell line H20961 was provided by the Gilad laboratory (Gallego Romero et al., 2015), and hiPS cell line 6032-4 was previously described (Yoon et al., 2019). Cultures were tested for and maintained Mycoplasma free. Lines were routinely propagated feeder-free in mTeSR1 medium (StemCell Technologies) on cell culture plastics coated with Geltrex basement membrane matrix (Gibco). Differentiation into hCO was performed under feeder-free conditions as previously described (Yoon et al., 2019).

*Generation of isogenic hiPS cell lines*

Guide RNAs (gRNAs) were designed to target the 30-mer repeat sequence (Rep-gRNA-9 and Rep-gRNA-12 in Table S5) and *in vitro* transcribed as previously described (Wucherpfennig et al., 2019). 0.5-1 $\mu$L of 40 $\mu$M Cas9-NLS purified protein (QB3, UC Berkeley) was complexed with 1 $\mu$g of each sgRNA for 5 minutes at room temperature. This complex was

*Corresponding authors
[1]These authors contributed equally to this work

nucleofected into $2x10^5$ cells along with 1 $\mu$l of 100 $\mu$M of a single-stranded oligo donor (ssODN) containing the single 30 bp repeat sequence found in chimpanzees and short homology arms (PtRep-ssODN in Table S5). Nucleofection was done with the P3 Primary Cell 96-well Nucleofector Kit (Lonza) on the Amaxa 96-well Shuttle Device (Lonza) using program CA-137 for 6032-4 and program DN-100 for H20961. The appropriate program for each cell line was determined by nucleofecting pMAX-GFP (Lonza) using each of the recommended nucleofection programs provided by the manufacturer. Cells were then assessed after 2 days for nucleofection efficiency and cell viability.

Following nucleofection, cells were allowed to recover in 1 well of a 96-well plate for 2 days and were then seeded at $1 - 5x10^3$ cells in a 10-cm plate. Colonies were picked using the EVOS FL microscope (Thermo Fisher) into a 96-well plate. When colonies were confluent to split, they were split 1:4 into a new 96-well plate. The remaining cells were resuspended in TE buffer, heated at 95°C for 10 min to lyse the cells, incubated at 55°C for 1 h with Proteinase K (Thermo Fisher, cat #25530015), and then heated at 95°C for 10 min to denature the Proteinase K. 1/20 of each sample was used for genotyping with primers Rep-sgRNA-TF and Rep-sgRNA-TR (Table S5). Lesions were identified on an agarose gel and bands that matched the size of the target lesion were extracted (Qiagen, cat #28704) for Sanger sequencing. The alleles in promising clones were then confirmed by cloning the gel extraction into the TOPO TA vector and performing colony PCR followed by Sanger sequencing. hiPS cell clones were then expanded for further use.

If needed, a second round of nucleofection was performed to make the locus homozygous using gRNAs targeting the incomplete lesion. The target lesion was further confirmed by Southern blot as previously described (Song et al., 2018) and by using primers that were multiple kilobases away from the gRNA target sites (Rep-sgRNA-TF2 and Rep-sgRNA-TR2 in Table S5) to ensure that the appearance of homozygosity was not due to lesions in the primer locations of Rep-sgRNA-TF and Rep-sgRNA-TR. Wild-type control clones were produced by nucleofecting cell lines with Cas9-NLS and PtRep-ssODN only and expanding single clones as described above. Wild-type control clones were referred to as Hs-TRACT$^{+/+}$, and hiPS cell lines where TRACT was replaced with the lone 30 bp sequence found in chimpanzees were referred to as Hs-TRACT$^{-/-}$.

*Fura-2 calcium imaging*

Dissociated telencephalon derived from Mm-hTRACT$^{-/-}$ x Mm-hTRACT$^{-/-}$ or Mm-hTRACT$^{KI/KI}$ x Mm-hTRACT$^{KI/KI}$ matings at E15.5 were cultured on poly-L-ornithine and laminin (Sigma) coated coverslips for 7-10 days. All embryos in each litter were pooled before plating. Dissociated Hs-TRACT$^{+/+}$ or Hs-TRACT$^{-/-}$ cortical organoids at D188-254 were cultured on poly-L-ornithine and laminin (Sigma) coated coverslips for 20-30 days. Mouse and human cultures were infected with AAV-DJ capsids encoding the eYFP reporter under the control of the human Synapsin-1 (Syn1) promoter to label mature neurons. Human neural cultures were also labeled with a Dlxi1/2b reporter to exclude GABAergic neurons from analysis. The cultures were incubated with 1 $\mu$M Fura-2 acetoxymethyl ester (Fura-2AM; Invitrogen) for 25 min at 37°C in NM medium (1X B27, 2 mM Glutamax, and 100 U/ml Penicillin/Streptomycin in Neurobasal Medium [Gibco, cat #21103049]), washed for 5 min and placed in a perfusion chamber on the stage of an inverted fluorescence microscope (TE2000U; Nikon) in low-KCl Tyrode's solution (5 mM KCl, 129 mM NaCl, 2 mM $CaCl_2$, 1 mM $MgCl_2$, 30 mM glucose, 25 mM HEPES, pH 7.4). Cells were then stimulated with high-KCl Tyrode's solution (67 mM KCl, 67 mM NaCl, 2mM $CaCl_2$, 1 mM $MgCl_2$, 30 mM glucose and 25 mM HEPES, pH 7.4) without or with nimodipine (final concentration 5 $\mu$M). Imaging was performed at room temperature (25°C) on an epifluorescence microscope equipped with an excitation filter wheel and an automated stage. Openlab software (PerkinElmer) was used to collect and quantify time-lapse excitation ratio images. Fluorescence images of hSYN1+ neurons were analyzed using the IGOR Pro software (WaveMetrics) to determine the peak amplitude.

*RNA extraction of mouse brain*

Mouse samples were dissected on ice and placed immediately in RNAlater (Thermo Fisher, cat #AM7021). Samples were stored overnight at 4°C and then stored for up to 4 months at -20°C. Prior to RNA extraction, samples were removed from RNAlater, placed into Trizol, and homogenized on the FastPrep-24 (MP Biomedicals) using Lysing Matrix D 2 ml tubes (MP Biomedicals, cat #116913100). For embryonic samples, the homogenate was directly applied to the Direct-Zol RNA Miniprep kit (Zymo Research, cat #R2051) for RNA extraction per the manufacturer's instructions. For post-natal samples, RNA was extracted from Trizol as previously described (Rio et al., 2010). 40 $\mu$g of RNA was then subject to DNaseI digestion to remove DNA contamination and applied to the Direct-Zol RNA Miniprep kit for clean-up.

*RNA extraction of hCO*

hCO samples were flash frozen and stored at -80°C as a pellet. Multiple organoids were collected and pooled for each sample. Prior to RNA extraction, samples were resuspended in either Qiagen buffer RLT or Trizol (Invitrogen, cat #15596018) and homogenized on the FastPrep-24 using Lysing Matrix D 2 ml tubes (MP Biomedicals). The homogenate was directly applied to the RNeasy Mini Kit (Qiagen, cat #74106) or the Direct-Zol RNA Miniprep kit (Zymo Research, cat #R2051) for RNA extraction per the manufacturer's instructions.

*RNA extraction of cultured cells*

Dissociated telencephalon derived from Mm-hTRACT$^{-/-}$ x Mm-hTRACT$^{-/-}$ or Mm-hTRACT$^{KI/KI}$ x Mm-hTRACT$^{KI/KI}$ matings at E15.5 were cultured on poly-L-ornithine and laminin (Sigma) coated coverslips for 7-10 days. All embryos in each litter were pooled before plating. Dissociated Hs-TRACT$^{+/+}$ or Hs-TRACT$^{-/-}$ cortical organoids at D160 were cultured on poly-L-ornithine and laminin (Sigma) coated coverslips for 20-30 days. Cells were stimulated with high-KCl Tyrode's solution (67 mM KCl, 67 mM NaCl, 2mM mM $CaCl_2$, 1 mM $MgCl_2$,

30 mM glucose and 25 mM HEPES, pH 7.4) for 1 h or 6 h. Control cells were processed concurrently through exposure to low-KCl Tyrode's solution (5 mM KCl, 129 mM NaCl, 2 mM $CaCl_2$, 1 mM $MgCl_2$, 30 mM glucose, 25 mM HEPES, pH 7.4) for 1 h or 6 h. Cells were collected in RDT buffer with beta-mercaptoethanol and processed using Qiagen RNeasy Micro kit (cat #74004) for RNA extraction per the manufacturer's instructions.

*Library preparation for RNA sequencing*

RNA sequencing libraries were prepared with the TruSeq Stranded mRNA Library Prep (Illumina, cat #20020595) using the IDT for Illumina – TruSeq RNA UD Indexes (96 Indexes, 96 Samples) (Illumina, cat #20022371) according to the manufacturer's instructions with one modification. Prior to PCR amplification, 10% of a subset of samples were run under the recommended PCR conditions with SybrGreen on the QuantStudio 5 qPCR machine (Thermo Fisher). We identified the number of PCR cycles required to reach the crossing point by qPCR and used that number of cycles for PCR amplification on the entire set of samples. We ran 8-10 PCR cycles for mouse samples and 10-11 PCR cycles for human samples. RNA quality was assayed with the Bioanalyzer 2100 (Agilent), and only samples with RIN > 8 were used for RNA sequencing. Library quality was assessed on the Bioanalyzer 2100, and library concentration was measured by qubit. Libraries were pooled and sequenced on the HiSeq 4000 (Novogene).

*CACNA1C total RNA levels and allele-specific expression*

*Cacna1c* was not differentially expressed in dorsal and ventral telencephalon at E12.5 and E15.5 or in the adult brain between Mm-hTRACT$^{-/-}$ and Mm-hTRACT$^{KI/KI}$ mice (Fig. S1C, E). In the Mm-hTRACT$^{-/-}$ and Mm-hTRACT$^{KI/KI}$ mice generated in the C57Bl6 inbred background, no differentially expressed genes were observed within 2 Mb of TRACT at E12.5 or E15.5 (Fig. S1E). *CACNA1C* and nearby genes were also not differentially expressed in hCO differentiated from Hs-TRACT$^{+/+}$ and Hs-TRACT$^{-/-}$ hiPS cell lines at D24, D50, and D100 (Fig. S4; Table S3). There were also no differences in the expression of *CACNA1C* or nearby genes in the unstimulated, dissociated mouse or human neurons (Table S2).

To profile differences in *Cacna1c* expression across additional brain regions and timepoints, we took advantage of the Mm-hTRACT$^{-/-}$ and Mm-hTRACT$^{KI/KI}$ mice generated on the mixed 126S6/C57Bl6 background where hTRACT was inserted on the 126S6 allele. We examined allele-specific *Cacna1c* expression with pyrosequencing in the dorsal telencephalon, ventral telencephalon, diencephalon, mesencephalon, and rhombencephalon at E16.5 and the cerebral cortex, hippocampus, midbrain/pons/medulla, cerebellum, and all other subcortical structures in P3, 4-week, 8-week, and 14-week mice.

Pyrosequencing assays were developed for rs30558791 (chr6:118696326, mm10) and rs30899647 (chr6:118627288, mm10). Rs30558791 is located in exon 13 of *Cacna1c* and has the genotype T in C57Bl/6 inbred mice and G in 129 inbred mice. Rs30899647 is located in exon 32 of *Cacna1c* and has the genotype C in C57Bl/6 inbred mice and T in 129 inbred mice. Genotypes were confirmed by Sanger sequencing in wild-type C57Bl/6 and 129 mice and in the repeat knock-in mouse lines with the primers rs30558791-F and rs30558791-R for rs30558791 and the primers rs30899647-F and rs30899647-R for rs30899647 (Table S5).

Pyrosequencing assays were developed using PyroMark Assay Design 2.0 (Qiagen). For rs30558791, we used the forward primer rs30558791-pyroseq-F, the reverse primer rs30558791-pyroseq-R (biotinylated at the 5' end and HPLC purified), and the sequencing primer rs30558791-pyroseq-S (Table S5). For rs30899647, we used the forward primer rs30899647-pyroseq-F (biotinylated at the 5' end and HPLC purified), rs30899647-pyroseq-R, and the sequencing primer rs30899647-pyroseq-S (Table S5). To verify the assays, we cloned the C57Bl/6 and 129 alleles into the TOPO TA vector (Invitrogen, cat #451641) to use as controls and mixed the two plasmids in ratios from 1:3 to 3:1 to a final concentration of 0.1 ng/$\mu$l. We confirmed that both assays correctly detected the proportion of each allele.

RNA was extracted as described above, and cDNA was made using the SuperScript VILO cDNA Synthesis Kit (Invitrogen, cat #11754250). cDNA was diluted 1:10 or 1:20 and amplified for 45 PCR cycles with DreamTaq Master Mix (Thermo Scientific, cat #K1072). Pyrosequencing was performed on the PyroMark Q24 (Qiagen) per the manufacturer's instructions, and data was analyzed with the PyroMark Q24 2.0.6 software (Qiagen). Results were concordant between rs30558791 and rs30899647. Results for rs30558791 are shown in Fig. S5. We did not observe a significant difference between the allelic ratios of *Cacna1c* expression between Mm-hTRACT$^{-}$ and Mm-hTRACT$^{KI}$ alleles at any time point or brain region assayed.

*Library preparation for isoform sequencing (Iso-Seq)*

Samples for isoform sequencing were pools of 2-3 biological replicates of the same genotype and species (Table S5). 400 ng of RNA, except for sample L2_PL6 for which only 300 ng was available, were reverse transcribed using GoScript RTase (Promega, cat #A2791) with oligo dT priming according to the manufacturer's protocol. For mouse samples, full length *Cacna1c* transcripts were amplified using cDNA template equivalent to 100 ng RNA, the mouse forward and reverse primers (Table S5), and PrimeStar GXL (Takara, cat #R050A). Temperature cycles of 98°C for 10 seconds (s), 57°C for 15 s, and 68°C 7 min, plus an initial denaturation of 98°C for 2 min and ending with an additional 68°C for 5 min, were performed for 35 cycles for the E15.5 samples and 32 cycles for the P2 samples.

Due to the low abundance of *CACNA1C* mRNA in the hCO samples and the reduction in PCR efficiency caused by the barcode adaptor sequences, *CACNA1C* transcripts were amplified from the human samples using a two-step PCR protocol. The first PCR amplified *CACNA1C* with primers containing only the *CACNA1C*-specific sequence, then the products were used as the template in the second PCR that added the barcode adaptor sequences. For the first PCR, full length *CACNA1C* transcripts were amplified from the human samples using cDNA template equivalent to 100 ng RNA, the human PCR 1 forward and reverse primers (Table S5), and PrimeStar GXL. 20 PCR cycles of 98°C for 10 s, 57°C for 15 s, and 68°C for 7 min, plus an initial denaturation of 98°C for 2 min and ending with an additional 68°C for 5 min, were performed. Excess primers were removed from the PCR products using the Monarch PCR Clean Up kit (NEB, cat #T1030S). The products were eluted in 15 $\mu$l elution buffer and added as template to the second PCR. The second PCR used PrimeStar GXL and the human PCR 2 forward and reverse primers (Table S5), and was performed for 12 cycles of the same temperature cycle as the first PCR.

The ~6.5 kb PCR products from both mouse and human samples were then each gel purified using a 1% agarose/TAE gel with GelGreen (Biotium, cat #41005) and the Monarch DNA Gel Extraction Kit (NEB, cat #T1020L). Product concentration was determined using the Broad Range DNA assay (Qubit) and 30 ng DNA was added to each barcoding PCR. The barcoding PCR used PrimeStar GXL and the EXP-PBC096 PCR Barcoding kit (ONT), and was performed for 15 cycles of 98°C for 10 s, 62°C for 15 s, and 68°C for 7 min, plus an initial denaturation of 98°C for 2 min and ending with an additional 68°C for 5 min. The products were purified using the Monarch PCR Clean Up kit and the DNA product molecular weight was checked using gDNA ScreenTape (Agilent). The concentration of each barcoded product was determined using the Broad Range DNA assay (Qubit).

The barcoded products were pooled in equal amounts, and the pool was 0.4X bead purified to remove any low molecular weight DNA using Mag-Bind Total Pure NGS beads (Omega Bio-tek, cat #M1378). The concentration and DNA size distribution of the purified pool was determined using the Broad Range DNA assay (Qubit) and gDNA ScreenTape (Agilent) respectively. 1$\mu$l of the purified pool of DNA was prepared using the SQK-LSK110 Ligation Sequencing kit (ONT) and was sequenced on a FLO-MIN106 flow cell (ONT) for 72 h, generating 2.2 million reads.

*$Ca_v$1.2 protein modeling*

To investigate isoform-specific differences in amino acid interactions within the $Ca_v$1.2 voltage-gated calcium channel, we used a recently resolved high-resolution structure of $Ca_v$1.2 (PDB: 8EOI) as a model (Chen et al., 2023). Using PyMOL, we modified exon 32 of the $Ca_v$1.2 structure to exon 31 to generate an alternate isoform.

Following exon modification, energy minimization was performed to relax the modified structure and ensure that it was in a stable conformation. Minimization was conducted using PyMOL's integrated molecular mechanics tools, which optimize bond lengths, angles, and dihedral angles by iteratively adjusting atomic positions to achieve a local energy minimum.

To assess changes in amino acid interactions between $Ca_v$1.2 containing exon 32 and $Ca_v$1.2 containing exon 31, we used PyMOL's distance and clash measurement tools, focusing on key residues in and around the modified exon region. We analyzed differences in hydrogen bonds and other non-covalent interactions to determine if the exon switch altered the local or global structural integrity or stability of the protein.

*Calcium imaging analysis*

Openlab software (PerkinElmer) and IGOR Pro (v.5.1, WaveMetrics) were used to collect and quantify time-lapse excitation 340:380 ratio images at an imaging rate of approximately 1 Hz. Peak amplitude was calculated by determining the maximum value post stimulation and subtracting the mean value of pre-stimulation baseline from it. For the traces, a representative field of view was selected for each experiment group, and the mean and the standard error of means were plotted for all recorded cells in that view. When put on the same plot, the baselines of different experiment groups were manually aligned. For the dot plots, significance was tested using Kruskal–Wallis test and Dunn's multiple comparison test.

*RNA sequencing analysis*

Sequencing reads were trimmed for adapter sequences using cutadapt v1.8.1 (Martin, 2011), and read quality was confirmed using fastqc v0.11.9 (Andrews et al., 2010). Reads were aligned using STAR v2.7 (Dobin et al., 2013) with two-pass mapping to either the mm10 or hg38 genome. The number of reads that overlap each gene was counted with featurecounts from the subread v1.6.0 package (Liao et al., 2014) using the Gencode vM24 annotation for mouse samples and the Gencode v29 annotation for human samples (Frankish et al., 2019). One library made from mouse neurons without TRACT at 6 h KCl was low quality (outlier in percent of overrepresented sequences and mapping percentage) and was excluded from further analysis.

Differential gene expression analysis was then performed with DESeq2 (Love et al., 2014) using default parameters. Embryonic samples were collected from both male and female mice, whereas adult samples were collected only from male mice. To adjust for differences in litter, sex, and dissections, we identified surrogate variables (SVs) representing batch effects using the sva package (Leek et al., 2012). We included 3 SVs as covariates for both adult and embryonic mice. 3 SVs produced a normal p-value distribution and minimized the number of gene expression differences between wild-type mice from the two lines. No SVs were included when analyzing hCO or activity-dependent transcription in mouse or human neurons.

Principal component analysis was performed using DESeq2. Gene ontology enrichment analysis was performed using clusterProfiler (Yu et al., 2012) with the background set at all expressed genes (genes with at least 5 reads in 50% of samples and could be analyzed by DESeq2). Genes associated with intellectual disability (ID) were downloaded from SysNDD on 4/15/2025 (Kochinke et al., 2016). Genes associated with autism spectrum disorder (ASD) were downloaded from SFARI (Abrahams et al., 2013) on 9/4/2024. Genes associated with epilepsy were downloaded from GeneTrek (Leblond et al., 2021) on 9/4/2024 using the gene list "High Confidence Epilepsy Genes". Significant enrichment of genes with concordant changes in activity-dependent expression with TRACT compared to all expressed genes was performed with the hypergeometric test with the Benjamini-Hochberg correction. We visualized only high-confidence SFARI genes (score = 1), but the results are similar when examining all SFARI genes.

Schizophrenia (SCZ) GWAS loci are from Table S2 in (Trubetskoy et al., 2022), and bipolar disorder (BPD) GWAS loci are from Table S2 in (Mullins et al., 2021). Significant enrichment near GWAS loci of genes with concordant changes with TRACT was performed using the "gene regulatory domain" parameters from default GREAT (McLean et al., 2010) and the binomial test with the Benjamini-Hochberg correction. Enrichment was performed relative to all expressed genes. Data visualizations were generated using ggplot2 (Wickham, 2016), pheatmap (Kolde, 2015), and karyoploteR (Gel and Serra, 2017).

The differential regulation index, DRI(condition1 / condition2), was calculated as $log_2(FC_{condition1}$(expression with stimulation / expression without stimulation) / $FC_{condition2}$(expression with stimulation / expression without stimulation)), where condition1 was either control human samples or samples with TRACT and condition2 was either wild-type mouse samples or samples without TRACT, respectively. Genes with concordant changes are activity-dependent genes (significant difference in gene expression with and without KCl stimulation in mouse or human neurons, with or without TRACT) and that have the same sign of DRI across models. Activity-dependent fold changes in mouse and human neurons after 4 h KCl from (Qiu et al., 2016) were downloaded from the manuscript. Processed read counts after 6 h KCl from (Ataman et al., 2016) were provided by the Greenberg lab. To compare across our dataset and previously published datasets, we restricted analysis to late response genes (activity-dependent in mouse or human neurons, with or without TRACT after 6 h KCl stimulation, in our dataset) that were analyzed in all three datasets.

*Pyrosequencing analysis*

Each sample was run in triplicate, and control plasmids were included in each sequencing run. Statistical significance was assessed with the Wilcoxon rank-sum test and corrected for multiple hypothesis testing at each time point.

*Iso-Seq analysis*

Human and mouse genomes (hg38, GRCm39 respectively) and annotations (GENCODE v35, GENCODE vM27 respectively) were downloaded from GENCODE (https://www.gencodegenes.org/). Base calling was performed with guppy v4.0.11 (ONT) and the *CACNA1C* transcripts were identified, annotated and quantified using the TAQLoRe pipeline (Clark et al., 2020). Briefly, reads passing QC and with an identified barcode were mapped to their respective transcriptomes using LAST v926 (Kiełbasa et al., 2011). Reads with a minimum 50% of their length mapping and supporting at least 80% of the transcript were retained to identify potential novel splicing events characterized by a minimum insert length of nine nucleotides. Novel events were confirmed through mapping of the reads to their respective genomes enabling the identification of their genomic coordinates. Supported novel exons were then incorporated into the *CACNA1C* metagene model prior to remapping the reads using GMAP v2019-03-15 (Wu and Watanabe, 2005) to enable transcript annotation, and quantification of relative expression. Novel exons were filtered by parsing the mapping CIGAR strings and retaining only exons with minimum length of 6 nucleotides and at least 70% support from the sequencing reads. All transcripts were checked and retained if they possessed at least 1 read per sample and found to be expressed in at least 2 libraries and with a minimum of 100 reads across all libraries following (Clark et al., 2020). This retained 67 exons in mouse samples and 76 exons in human samples. Transcript quantification was tmm-normalized across all samples. The proportion of isoforms were then normalized within each sample to account for differences in sequencing depth. Unfortunately, one sample, RW_L1, had very low sequencing counts, so RW_L1 and its paired sample RK_L1 were excluded from downstream analysis.

To compare isoform usage across species, orthologous mouse and human exons were identified. 55 of the 67 mouse exons were assigned to 1+ orthologous human exons, and 64 of the 76 human exons were assigned to 1+ orthologous mouse exons. In addition, 47 mouse isoforms map to 60 human isoforms. Exons that do not have an orthologous match were lowly expressed and did not pass filtering in the other species or were excluded in the other species due to the location of the PCR primers. One-to-many mapping between mouse and human isoforms were due to differences in PCR primer location. The relative abundances of exons 31 and 32 between samples with TRACT and samples without TRACT were assessed using a two-sided t-test.

### Supplemental Figures

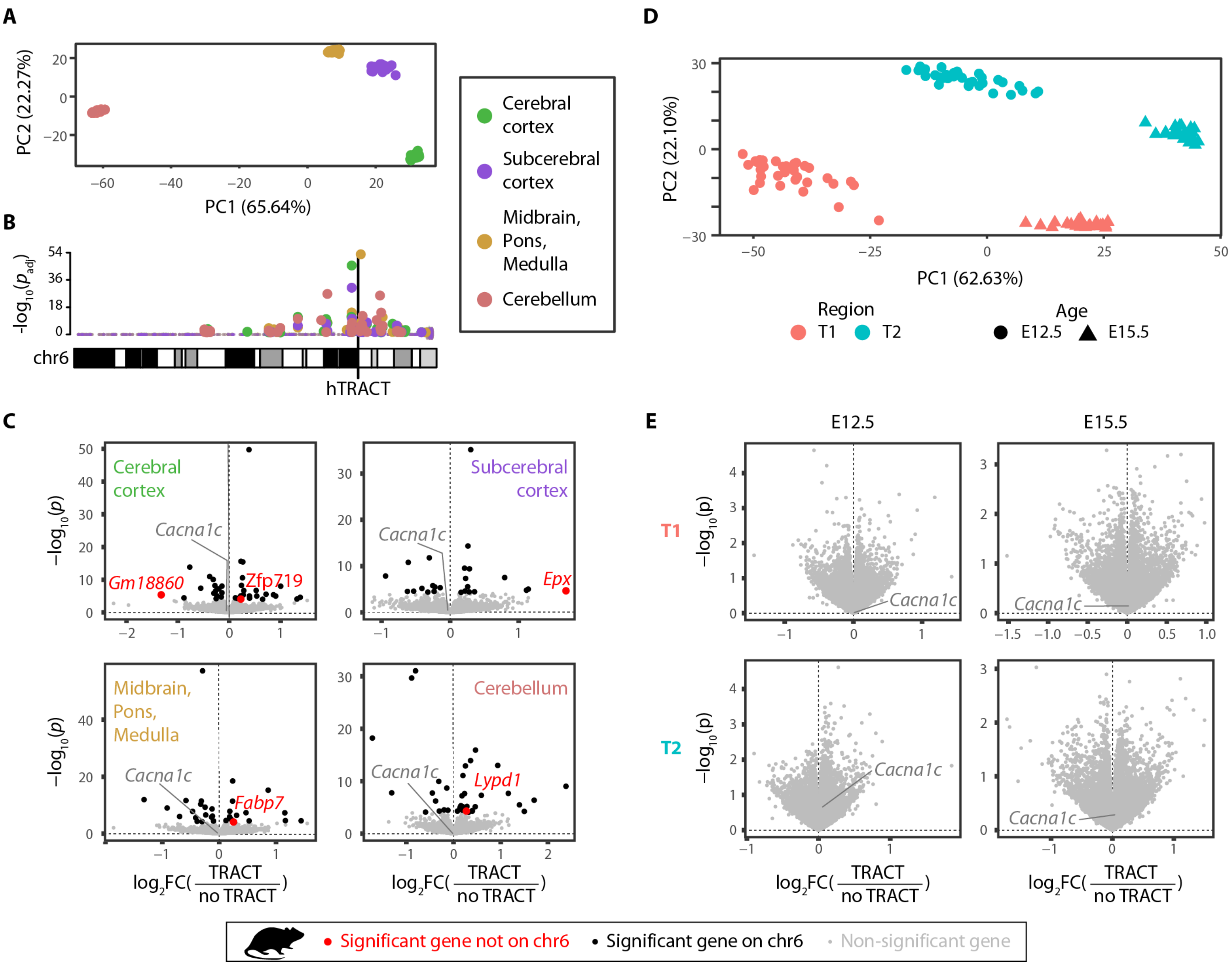


Figure S1: **Transcriptional effects of hTRACT on the adult and developing mouse brain.** (A) Principal component analysis (PCA) of RNA-seq of brain regions dissected from 14-week-old mice. (B) Differentially expressed genes between Mm-hTRACT$^{-/-}$ and Mm-hTRACT$^{KI/KI}$ cluster around the insertion site of hTRACT on chr6 for all brain regions. (C) Volcano plots of differentially expressed genes. Black: Differentially expressed genes on chr6. Red: Differentially expressed genes not on chr6. Gray: Non-significant genes. (D) PCA of RNA-seq of brain regions dissected from embryonic mice. (E) There are no differentially expressed genes between Mm-hTRACT$^{-/-}$ and Mm-hTRACT$^{KI/KI}$ in the dorsal telencephalon (T1) or ventral telencephalon (T2) at E12.5 or E15.5.

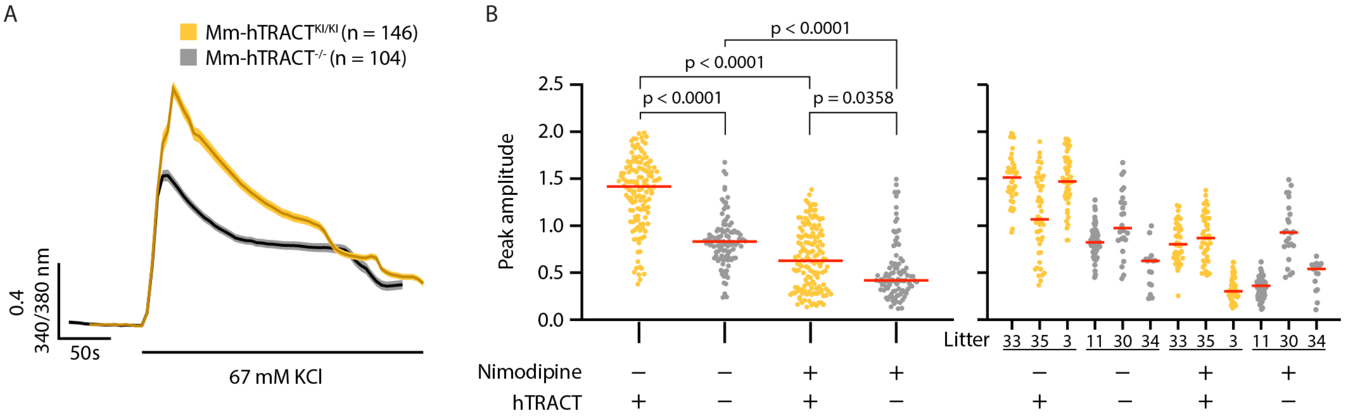


Figure S2: **Calcium response to KCl stimulation in neurons from a second mouse line.** (A) Representative traces of intracellular calcium levels following KCl depolarization from one field of view for neurons from a second mouse line (Materials and Methods). The y-axis represents the fluorescent intensity of the calcium reporter Fura-2. The lines and shaded areas indicate means and 95% confidence intervals, respectively. (B) Peak calcium amplitude with and without TRACT and/or nimodipine. Left, data pooled across mouse litters; right, data separated by litter. Each dot represents one cell (n = 479 cells); Kruskal–Wallis test, $p < 0.0001$. P-values of Dunn's multiple comparison test are labeled. Red bars show median values of each group.

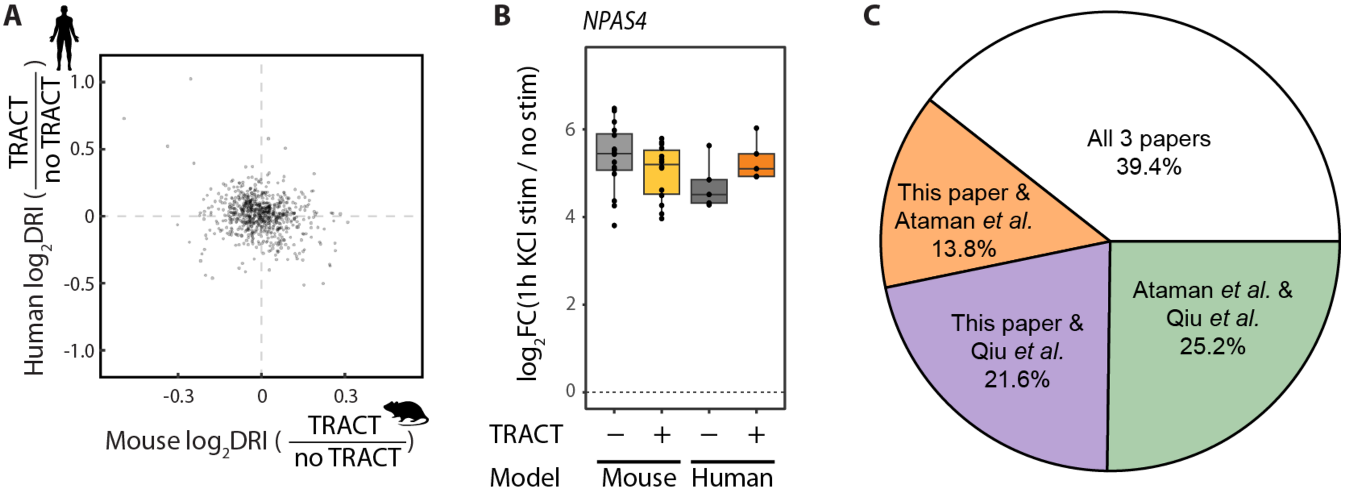


Figure S3: **Activity-dependent transcription after KCl stimulation.** (A) Differential regulation indices (DRI) for mouse and human are not correlated after 1 h of KCl stimulation. (B) The effect of TRACT on *NPAS4* expression is not concordant between mouse and human models after 1 h KCl. (C) Percentage of late response genes (4 h or 6 h KCl) with the same direction of effect on activity-dependent transcription (DRI(Hs/Mm) has the same sign) between (Qiu et al., 2016; Ataman et al., 2016), and this paper.

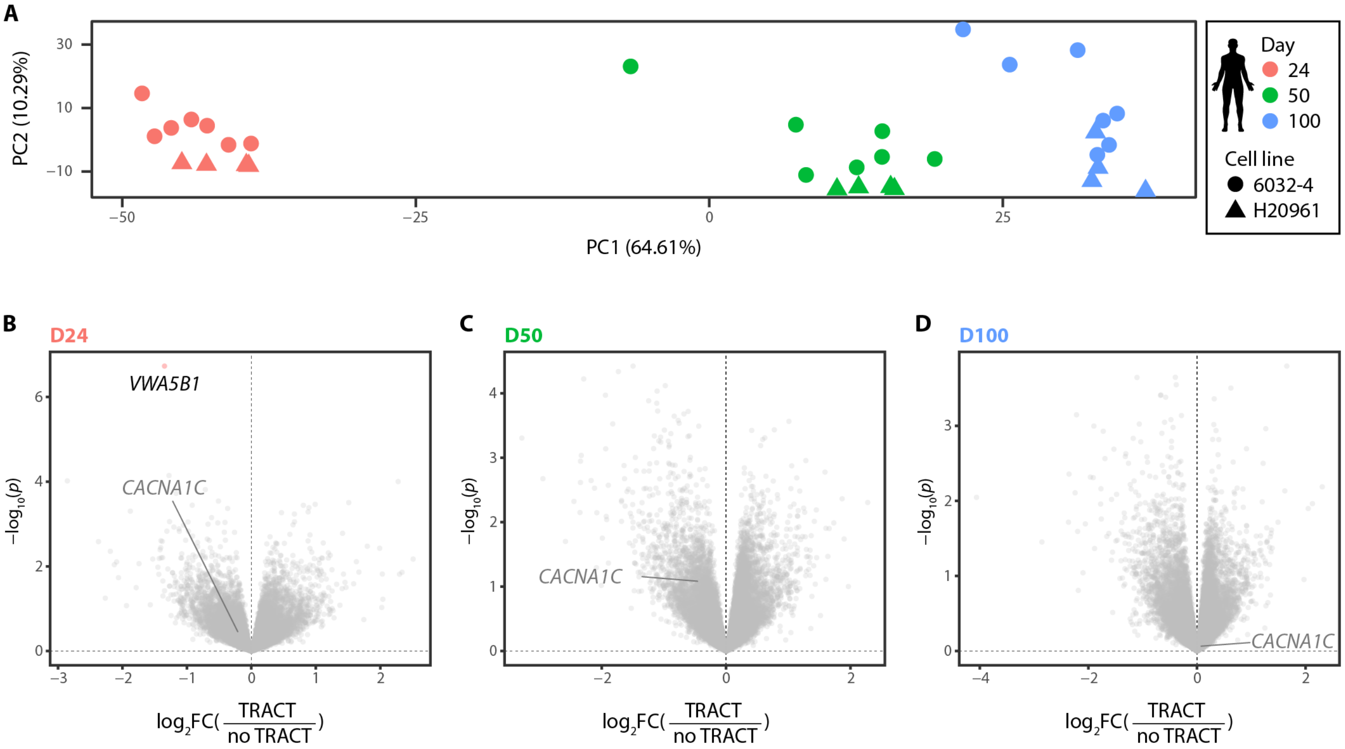


Figure S4: **Transcriptional effects of TRACT on hiPS cell-derived cortical organoids.** (A) PCA of RNA-seq of human cortical organoids at different time points. (B-D) Volcano plots comparing gene expression between Hs-TRACT$^{-/-}$ and Hs-TRACT$^{+/+}$ cortical organoids at each time point. *CACNA1C* itself is not differentially expressed at any assessed time point.

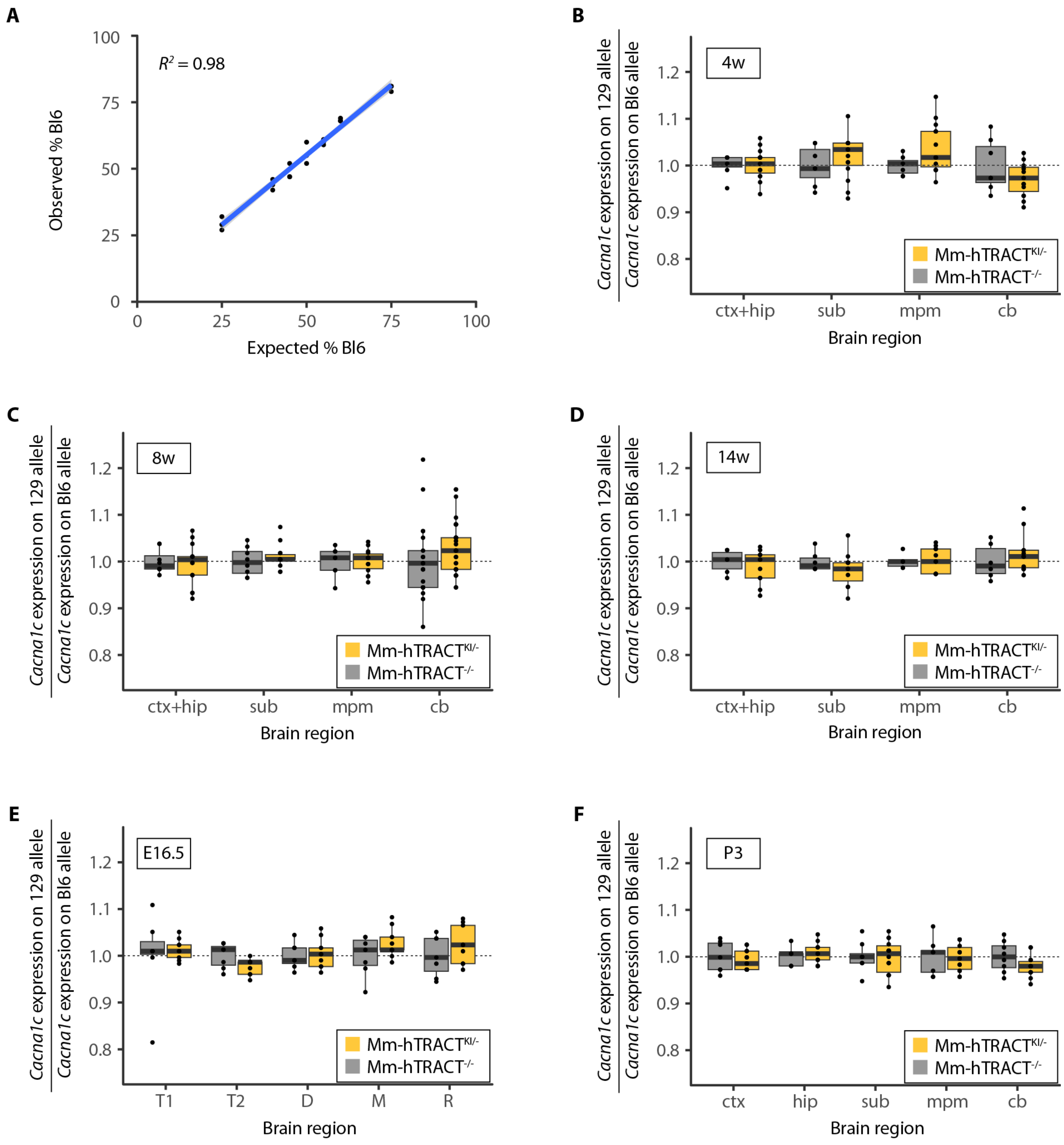


Figure S5: **Activity-dependent transcription after KCl stimulation.** Using pyrosequencing at rs30558791 (Materials and Methods), we compared *Cacna1c* expression between Bl6 and 129 alleles in Mm-hTRACT$^{KI/-}$ mice (yellow), where hTRACT is on the 129 allele and the Bl6 allele lacks hTRACT, and Mm-hTRACT$^{-/-}$ mice (gray), where both the 129 and Bl6 alleles lack hTRACT. (A) Calibration curve with standards containing known ratios of the Bl6 and 129 alleles at rs30558791. (B-F) Allele-specific expression of *Cacna1c* across different brain regions and developmental time points. hTRACT showed no consistent *cis*-effect on *Cacna1c* expression. Ctx: cerebral cortex. Hip: hippocampus. Sub: all other subcortical structures. Mpm: midbrain, pons, and medulla. Cb: cerebellum. T1: dorsal telencephalon. T2: ventral telencephalon. D: diencephalon. M: mesencephalon. R: rhombencephalon.

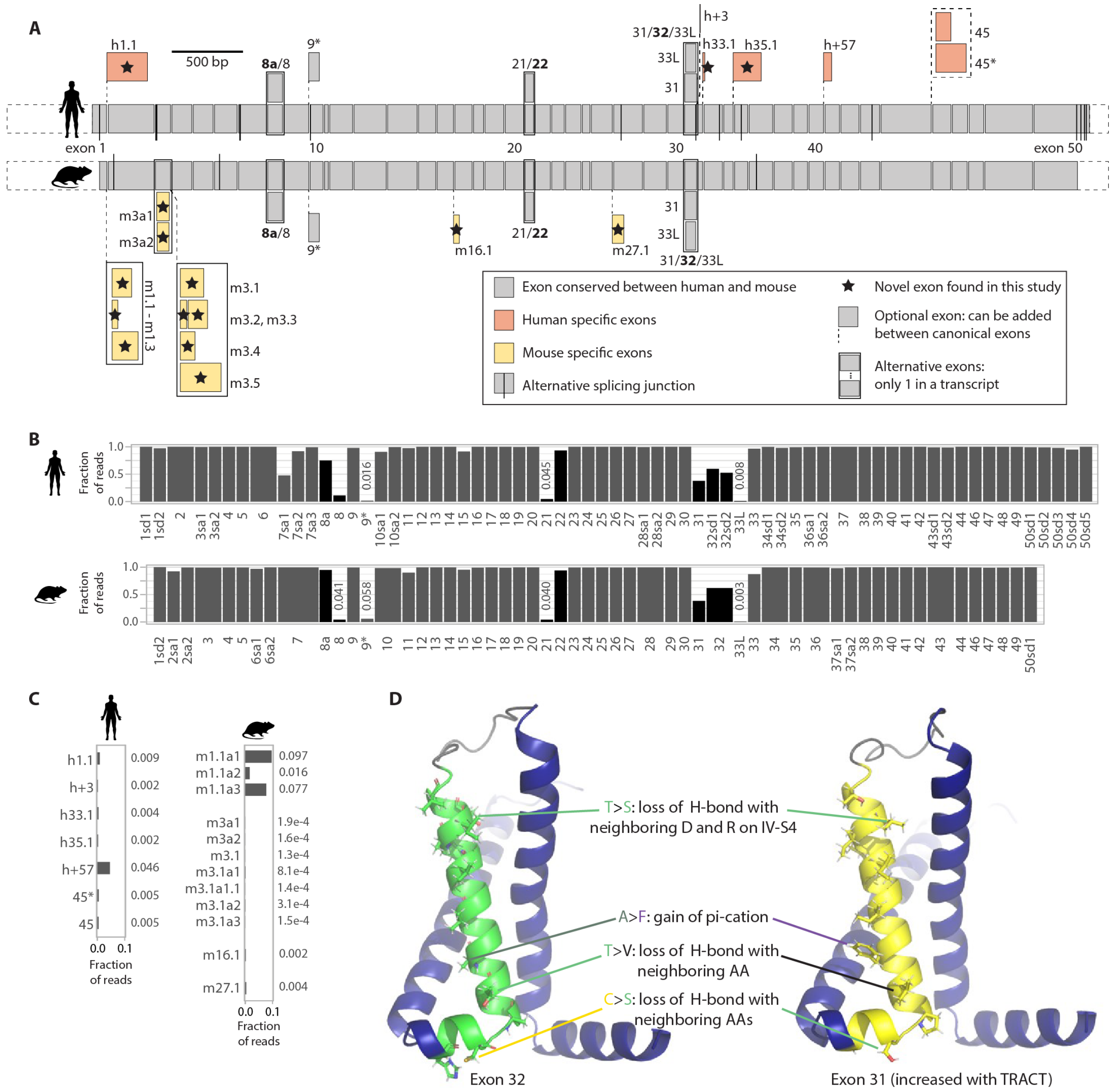


Figure S6: **Targeted long-read sequencing of *CACNA1C* isoforms.** (A) All exons and alternative splice sites in *CACNA1C* identified in this study for human (top) and mouse samples (bottom). Optional exons (shown with dotted lines) are exons that can be inserted between two exons. Alternative exons (shown stacked within a box) never appear together within the same transcript. For each alternative box, the dominant exon is labeled in bold. Optional alternative exons, e.g. m1.1 – m1.3, are mutually exclusive optional exons. m3.2 and m3.3 always co-occur. 45* and 45 were mutually exclusive except in one rare isoform. Note that exon 1 and exon 50 are actually much longer than shown here (indicated by the extending dashed lines). (B) Fraction of reads containing each exon in human (top) and mouse (bottom). Only exons identified in both species are shown. Exons with alternative splice sites are shown separately for each splice site and are labeled as sd1, sd2… or sa1, sa2… for alternative splice donor (sd) or splice acceptor (sa), respectively. Abundances of exons that appear in less than 10% of transcripts are labeled. (C) Proportion of reads containing each human- or mouse- specific exon. Note that the x-axis is between 0 and 0.1. (D) Structure of exon31 and exon32 with neighboring exons. Amino acid changes between exons 31 and 32 that are predicted to affect residue interactions are highlighted. For exons 31 and 32, N' to C' goes from the bottom to the top of the figure.

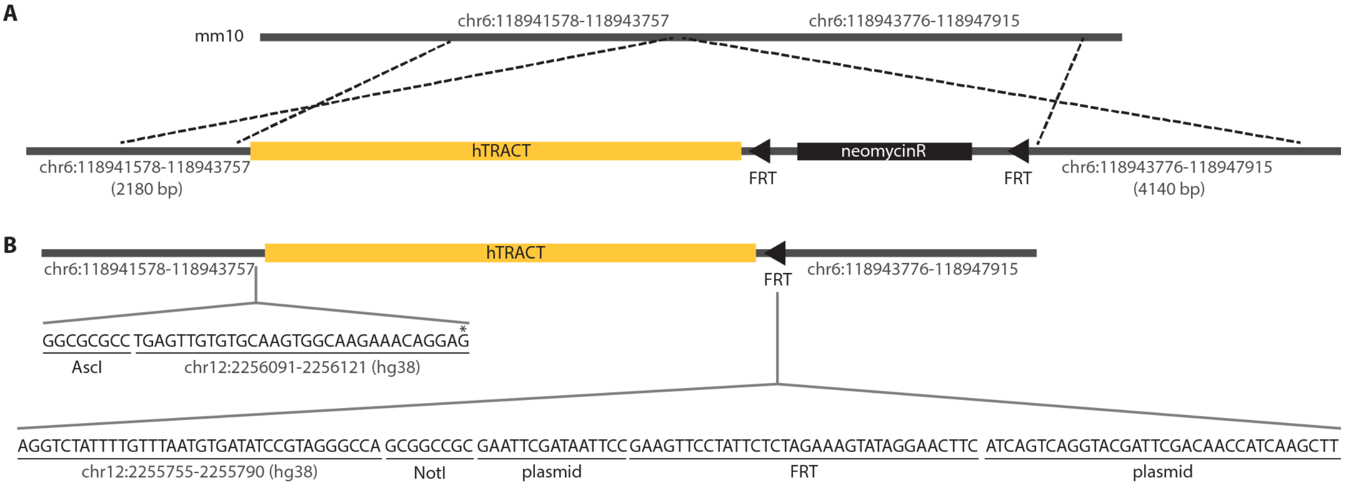


Figure S7: **Schematic of mouse knock-in lines.** (A) We generated two mouse knock-in lines with insertions of different hTRACT alleles found in humans (Materials and Methods). Each hTRACT allele was inserted into the same orthologous location in the mouse genome. The insertion sequence included the indicated homology arms and a neomycin resistance gene flanked by FRT sites. Knock-in mice were bred to mice expressing Flp recombinase to excise the neomycin resistance gene. (B) In addition to the inserted hTRACT alleles, the knock-in mouse lines also contain flanking human sequence on the 5' end (31 bp) and 3' end (36 bp), 2 restriction sites used for cloning (16 bp), a FRT site (34 bp), and flanking sequence from the plasmid backbone (49 bp). One of the knock-in mouse lines has an A at the position marked by an asterisk instead of a G.

**Supplemental Tables**

Table S1: Differential gene expression analysis between Mm-hTRACT$^{KI/KI}$ and Mm-hTRACT$^{-/-}$ mice
Table S2: Differential gene expression analysis after neuronal stimulation
Table S3: Differential gene expression analysis between Hs-TRACT$^{+/+}$ and Hs-TRACT$^{-/-}$
Table S4: Isoform sequencing
Table S5: Primers and oligos
